# Quantification of live cell membrane compartmentalization with high-speed single lipid tracking through interferometric Scattering Microscopy

**DOI:** 10.1101/2021.08.06.455401

**Authors:** Francesco Reina, Christian Eggeling, B. Christoffer Lagerholm

## Abstract

The specific details of the lateral diffusion dynamics in cellular plasma membrane are an open topic in modern biophysics. Many studies have documented several different behaviours, including free (Brownian) motion, confined diffusion, transiently confined (hop) diffusion, anomalous diffusion, and combinations thereof. Here we have employed Interferometric Scattering Microscopy (ISCAT) to explore the lateral diffusion dynamics in the plasma membrane of living cells of a biotinylated lipid analogue that had been labelled with streptavidin-coated gold nanoparticles (20 and 40nm in diameter) at a sampling rate of 2kHz. The data was analysed with an unbiased statistics-driven mean squared displacement analysis pipeline that was designed to identify both the most likely diffusion mode for a specific data set, and the best fit parameters of the most likely model. We found that the prevalent diffusion mode of the tracked lipids, independent of the particle size, is compartmentalized diffusion, although the use of the larger tags resulted in tighter confinement and reduced diffusion rates. Through our analysis and comparison with simulated data, we quantify significant physical parameters, such as average compartment size, dynamic localization uncertainty, and the diffusion rates. We hereby further demonstrate the use of a confinement strength metric that makes it possible to compare diffusivity measurements across techniques and experimental conditions.

**Statement of Significance:** This work offers new details on the data analysis of lipid diffusion on cellular membranes in vitro, through Interferometric Scattering microscopy. With this technique, we performed single particle tracking (SPT) experiments at 2kHz sampling rate. We analyzed the data through an unbiased statistics-driven protocol. The data shows that the diffusion motion of the tracked lipids follows mainly the “hopping” diffusion behaviour, whereby transient confinement zones hinder the particle dynamics. Matching the experimental data with diffusion simulations, we have been able to verify the physical parameters inferred by the experimental data analysis. Finally, we showcase a framework to compare SPT data with other techniques, to offer a complete overview of plasma membrane dynamics.

## Introduction

A dominant feature of lipid diffusion in the cellular plasma membrane, highlighted through single-particle tracking (SPT) (1–6) and super-resolution STED microscopy (6–8), is the presence of transient confinements (9,10). In compartmentalized diffusion, the molecules diffuse within compartments of a corralled surface, with a probability of “hopping” into a neighbouring compartment. This is compatible with the picket-fence model of the cellular plasma membrane, where the cortical actin meshwork, in interplay with linked picket-like transmembrane structures such as proteins, induces the compartmentalization (11). In fact, several possible mechanisms have been identified as the origin of such diffusion heterogeneities, such as structural properties of the cellular membranes (12,13), or the role of transmembrane proteins, such as CD44, in affixing the actin cytoskeleton to the plasma membrane (14). Resolving such characteristics requires both high spatial resolution, as compartment sizes are usually below 200 nm in size (15), and high temporal resolution for distinguishing intra-from inter-corral dynamics. This is only accessible to few experimental techniques (16–19), amongst which optical microscopy has proven particularly successful, and even more so through single molecule and super-resolution microscopy techniques. However, these sampling conditions introduces major challenges to observation techniques such as SPT, and explains why some details of this diffusion mode are still under debate (8,10).

Fluorescence-based SPT has the advantage of preserving the specificity characterizing fluorescence imaging while minimizing potential labelling artefacts. However, conventional fluorescence microscopy has not yet achieved framerates faster than 2kHz for studies on live cells (20,21). At the state of the art, however, the fastest sampling rates (up to 50kHz) have been reported in scattering detection-based experiments, through the use of larger nanoparticle tags and advanced camera equipment (22). Recent evolutions of Interference Reflection Microscopy (23,24), namely Interferometric Scattering (ISCAT) and Coherent Brightfield (COBRI) Microscopy, also managed to approach such levels of temporal resolution in SPT on cell membranes in recent years (25–30).

In ISCAT microscopy the reflected and backscattered originated by the coherent incident light generate an interference figure on the camera sensor (31). The imaging contrast is mainly given by the phase difference term between the incident and reflected field. This results in increased Signal to Noise (SNR) ratios compared to darkfield and brightfield microscopy (32). While this has allowed the use of smaller tags to probe particle motion, (25,33,34), the use of gold nanoparticles carries the obvious advantage of unparalleled scattering capabilities, leading to high signal-to-noise ratios. Although crosslinking is likely to happen (35), it has already be shown that it would only slow down the diffusion without altering the diffusion mode(36). Given the relative simplicity of the experimental setup and the data analysis, ISCAT has already been employed to perform SPT experiments on model and cellular membranes (25,26,42,27,28,36–41).

The analysis of SPT data is largely reliant on the relation between the Mean Squared Displacement (MSD), the diffusion coefficient (D) and the time interval at which the displacements are calculated *n δ_t_* (43–45). This relation does not account for inevitable experimental errors, such as the dynamic particle localization uncertainty, and motion blur effects that result from finite camera integration of continuously moving objects. In fact, it has already been shown that particle detection in the context of a finite localization uncertainty produces a positive offset to the MSD curve (46–48) while the effect of motion blur translates to a negative offset that is proportional to the time interval *δ_t_* (49). A collective consequence of these artefacts can be an underestimation of the localization uncertainty. The effect of this, in the case of slower sampling *(n δ_t_* ≥ 10 ms) where the actual displacement of each molecule between successive localizations is typically much larger than the localization uncertainty, is negligible (36). However, with faster sampling rates *(n δ_t_* ≤ 5 ms), where the actual displacement of each molecule between successive frames becomes comparable to (or even smaller than) the localization uncertainty, this effect is likely to cause misleading conclusions (6).

In this work, we employed ISCAT microscopy to detect single particle trajectories of biotinylated lipid analogues tagged with streptavidin-coated gold nanoparticles (Figure 1), in order to quantitatively describe the compartmentalization of the plasma membrane. To this end, we propose and validate a novel analysis pipeline to analyse single particle trajectories on plasma membrane, and to address the complexities thereof (10). One of the distinguishing features of this analysis pipeline is, first of all, the estimation of the dynamic localization uncertainty through the trajectories themselves. Further, we implement a statistics-driven method to classify the single particle trajectories amongst several plausible diffusion modes. Notably, we choose to compare the well-known anomalous diffusion model, against other models obtained from the analytical solution of particle diffusion in a corralled environment (50–52). Using our analysis pipeline, we show that the most suitable models to describe the diffusive motion of the tracked nanoparticle-tagged lipids are indeed these last models, describing compartmentalized (hop) diffusion, with a characteristic compartment size of ≈ 100 nm. We confirmed the accuracy of our conclusion by comparing the results obtained through our data analysis pipeline to simulated particle diffusion on a two-dimensional lattice. Finally, we adopt the confinement strength (S_conf_) metric, which allows straightforward comparisons between the present results with other related studies from the same or other related techniques. The use of this metric shows that the diffusive motion herein described appears to not be cell line-specific, as demonstrated by comparison with the data from past experiments (6,8).

**Figure 1.**
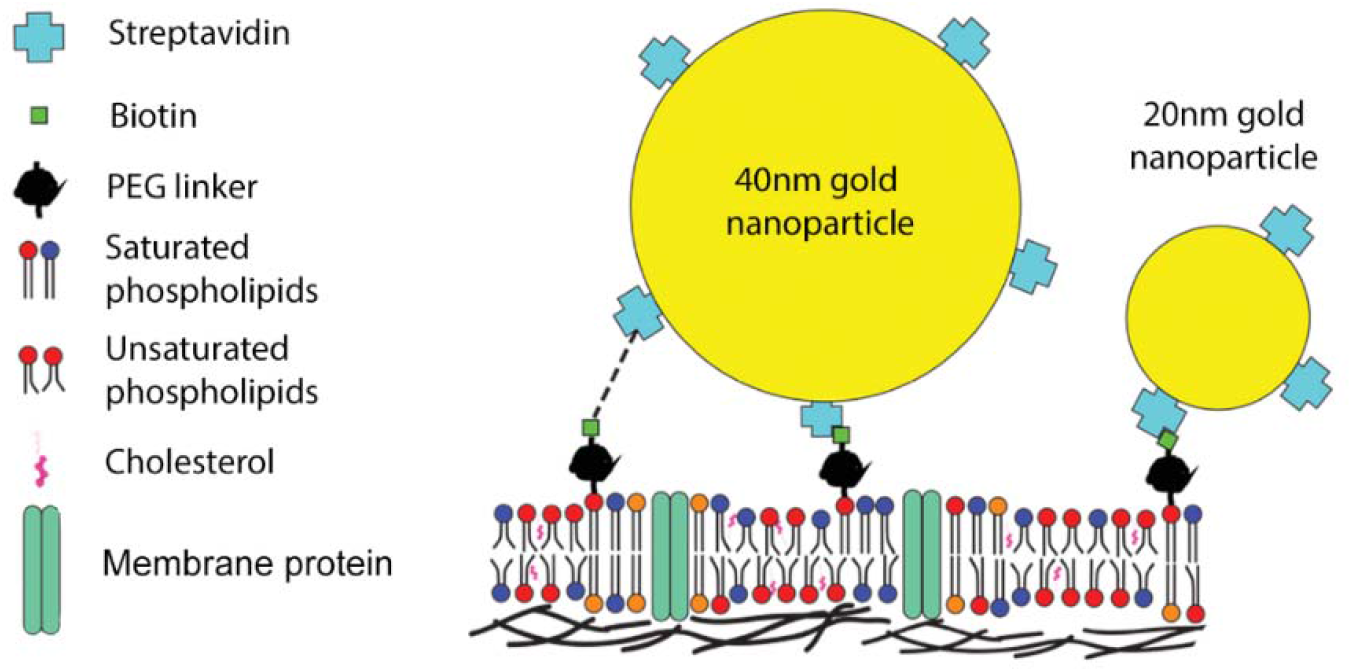
Scheme for the labelling of the cell surface. The scattering tags, gold nanoparticles of two different diameters (20nm and 40nm) target the biotinylated lipids inserted in the cellular membrane, owing to their streptavidin coating. The possible, although not certain, effects of cross linking are also highlighted (dashed line).

## Materials and Methods

### Lipids and cell line

Atto488-labelled DOPE(1,2-Dioleoyl-*sn*-glycero-3-phosphoethanolamine) was purchased from Atto-Tec. DSPE-PEG(2000)-Biotin (DSPE: 1,2-distearoyl-sn-glycero-3-phosphoethanolamine, 2kDa PEG linker between the phospholipid polar head and the biotin), henceforth referred to simply as DSPE-PEG-Biotin, was purchased from Avanti Polar Lipids. Lipid stock solutions were stored at −20C in chloroform. Gold Nanoparticles of 20 nm and 40 nm diameter (Ø), streptavidin coated, where purchased from BBI solution in stocks, the concentration of which is expressed as 10 OD (optical density). PtK2 cells derived from rat kangaroo (Potorus tridactylis) kidney tissue (53) were available in the laboratory. These were cultured following known protocols, growing them in Dulbecco Modified Eagle Serum (DMEM, Sigma Aldrich), supplemented with ~15% FBS (Fetal Bovine Serum), L-Glutamine, and Penicillin-Streptomycin (54). Before labelling and experiments, the cells were grown in sterile single-use flasks, placed in a 37C incubator in water vapour-saturated atmosphere with 5% CO2.

### Cell membrane labelling

PtK2 cells were seeded and left to proliferate on methylated-spirits cleaned glass supports (25mm diameter, #1.5 thickness coverslips), and used at a stage where they did not yet reach confluency. A sufficient separation between the cells is deemed necessary to ensure that the membrane of each cell was not affected by the presence of neighbouring cells that may cause deformation. This translates to an estimated 50-70% confluency. Before the labelling, to allow a more comfortable and secure application of the labelling protocol, the glass supports were mounted in a water-tight steel chamber (Attofluor chambers, Thermo Scientific). The cell labelling procedure was adapted from the protocol described in (55). A stock solution of DSPE-PEG-Biotin in 1:1 Chloroform-Methanol at 10mg/ml was desiccated via nitrogen gas flow, and the lipid suspended again in absolute ethanol to a concentration of 20 mg/ml. This was diluted in L15 medium to a final concentration of 0.2mg/ml, and incubated at 37C for 20-30 minutes. In the same buffer, a small concentration of Atto488-DOPE was dissolved, in order to facilitate detection of the labelled cells by using the fluorescent channel of the ISCAT microscope. After the incubation with the biotinylated lipids, the cells were washed with fresh L15 buffer, and incubated for 10-15 minutes at 37C with a solution of 0.6uM of streptavidin-coated 20nm or 40 nm diameter (Ø) gold nanoparticles in L15 buffer. Afterwards, the cells were once again rinsed with fresh L15 buffer, and used for the experiments. This protocol produced a sparse labelling of cells (~1-2 nanoparticles per cell, with multiple labelled cells).

### Interferometric Scattering and Total Internal Reflection Microscope setup

ISCAT experiments were performed on a custom built, following the protocol in (56), that has been previously described (36) with some useful modifications. The output from a 660nm solid-state laser diode (OdicForce) was scanned in two directions (equivalent to the x and y on the sample plane) by two acousto-optic deflectors (AOD, Gooch & Housego and AA Opto-Electronics). The scanned output was then linearly polarized, relayed to the back focal plane of the objective via a two-lens telecentric system, passed through a polarizing beam splitter and circularly polarized by a quarter wave plate (B.Halle). The light was finally focused by a Plan Apochromatic 60x, 1.42NA oil immersion objective (Olympus), mounted in an inverted geometry. As stated in the introduction, the reflected component by the glass-sample interface and the back-scattered component by the sample were collected by the same objective, and reflected onto the detection path by a polarizing beam-splitter. The final image is obtained by focusing these two interfering beams onto the CMOS camera sensor (Photonfocus MV-D1024-160-CL-8) to acquire time lapses with an effective magnification of 333x (31.8nm effective pixel size).

In addition to this imaging mode, the microscope was also equipped with a total internal reflection fluorescence-capable channel. A 462nm wavelength solid state laser diode output was focused on the back aperture of the objective. TIR illumination condition was achieved via a movable mirror, until the reflection of the illumination beam was visible on the other side of the back-aperture. The fluorescence signal was separated from this reflection by means of an appropriately placed dichroic mirror, and imaged onto a difference CMOS camera (PointGrey Grasshopper 3). The labelling of cells with a fluorescent lipid analogue ensured that the sample could be correctly identified in a second, independent way. However, this part of the setup was not optimized to perform fluorescence imaging experiments, and it was used merely as a guide for the user.

Stabilization of the imaging plane was achieved by a piezo-actuated objective positioner (PiezoSystem Jena) in open-loop configuration. This ensured enough stability in the focus to perform the intended measurements. A summarizing scheme for the imaging setup is given in Supplemental Figure S1.

### Imaging conditions

The glass support with the cells was positioned on the microscope stage while still inside the steel chamber used for labelling. The deformation in the support induced by the O-ring present in the steel chamber produced a drift in the apparent z position of the sample when changing area of imaging, but once readjusted, the sample was stable enough to allow prolonged observation times and correct recording, also thanks to the piezo-actuated objective positioner (MiPos, PiezoSystem Jena). The cells were imaged in L15 medium at a temperature of 37C, in room atmosphere and humidity, thanks to a temperature control dish (Warner Instruments). The laser power area density used to illuminate the cells for ISCAT imaging was 17.5 kW/cm2, and given the illumination wavelength used (660nm), temperature-induced artefacts can be ruled out (30). Although the aforementioned power density might seem high, it has been shown in similar cell lines that prolonged exposures to even higher power densities, at the same wavelength of our experiments, are well tolerated by this kind of sample (57). Using the CMOS camera previously mentioned, we collected 2000 frames long movies in a 200×200 px^2^ region of interest, with 0.227ms exposure time, resulting in 2kHz sampling rate and roughly 41 μm^2^ imaging area. Although similar samples and experiments have been carried out at much faster sampling rates, it has been shown that similar sets of parameters are sufficient to describe the scenario herein considered (3). An evaluation of localization precision with these conditions has been derived by measuring the FWHM of the distribution of relative distances of two immobilized gold nanoparticles on glass, according to the procedure in (26,58,59), giving the value of 2.6nm. More details on the estimation of the localization uncertainty are given in Supplemental Note 1.

### Trajectory detection

Single Particle Tracking data analysis requires the trajectories of the particles to be extracted from the collected movies. The movies were collected in TDMS file format with a LabView software (courtesy of the Kukura Laboratory, University of Oxford), and converted to TIFF image stacks with a home-written MatLab code, based on the ConvertTDMS function by Brad Humphreys (https://www.github.com/humphreysb/ConvertTDMS, last retrieved March 23, 2020). Before particle tracking, all the movies were elaborated by subtracting the median filter, as previously elucidated (36). In addition to this, the average intensity projection of the movie was obtained and subtracted from every movie frame, in order to separate the moving fraction of the sample from the static background (Supplemental Figure S2). Image processing was performed using the FIJI platform (60). Tracking was performed using the Spot Detection function in Imaris 9.5 (Bitplane, Oxford Instruments). Subsequent trajectories were imported in text format for post-processing in Mathematica (version 12.0.0.0; Wolfram Research) with custom written codes.

### Analysis of Single Particle Trajectories

We hereby give a short overview of the analysis pipeline for the analysis of single particle trajectories. For a more complete overview of the methodology adopted, see Supplemental Note 2. All the analysis routines herein described, can be easily reconstructed using the Python package reported in (61).

The protocol employed for this study is a refinement of that presented in (6,10). We calculate the apparent diffusion coefficient D_app_(t_n_) curve for each particle trajectory, defined as:

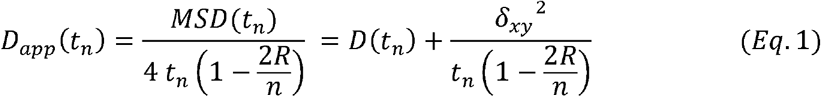

where MSD(t_n_) corresponds to the time-averaged mean squared displacement, t_n_ is the n-th time interval, defined as multiples of the time interval between successive localizations t_0_, R is the motion blur correction factor (6,49,62), D(t_n_) is the artefact-corrected time-dependent diffusion coefficient, and *δ_xy_* is the average dynamic localization uncertainty. In our workflow, we have set R = (1/6)(0.227ms/0.5ms) for full frame averaging, for imaging with a camera exposure time of 0.227ms with a frame rate of 2 kHz (49,62). We have furthermore restricted our analysis to truncated trajectory segments of 500 localizations each, to allow for constant statistical sampling of the D_app_ curves. Summary statistics of the number of raw trajectories, and the number of segments obtained from the truncation are reported in Table 1.

**Table 1.** Summary statistics for the trajectories analysed in this study. N_Trajectories_ refers to the total number of trajectories per sample, while 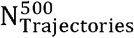 refers to the corresponding number of 500 localization-long segments extracted from the raw trajectories. In this instance, the gold nanoparticles are referred to as sAv-Au (Ø20nm) or sAv-Au (Ø40nm), as a reminder of the streptavidin (sAv) coating present on the probe.

| Sample | $N_{Trajectories}$ | Trajectory Length (±s.t.d.) | $N_{Trajectories}^{500}$ |
| --- | --- | --- | --- |
| DSPE -PEG2000-biotin / sAv-Au (∅20nm)<br>on Ptk2 cells | 92 | $1430 \pm 970$ | 229 |
| DSPE-PEG2000-Biotin / sAv-Au (∅40nm)<br>on Ptk2 cells | 165 | $1510 \pm 1040$ | 433 |
| sAv-Au (∅20nm) immobilized on glass | 4 | $4000 \pm 0$ | 32 |
| sAv-Au (∅40nm) immobilized on glass | 4 | $4000 \pm 1$ | 31 |

Our experimental data suffered from slight environmental vibration artefact with a frequency corresponding roughly to that of a common fan frequency of about 8500 rpm. This was most clearly visible in the control trajectory data of immobile gold particles on glass (Supplemental Figure S3). Prior to the quantitative analysis, we thus removed the influence of these environmental vibrations by median filtering in Fourier space the frequencies ranging from 140-160 Hz from the MSD curves of each single particle trajectory. A comparison between the original and the vibration corrected data is shown in Supplemental Figure S3, and the parameters for corrections are given in Supplemental Table S1.

By substituting the expressions of D(t_n_) shown in Table 2, corresponding to different plausible diffusive motion models (10) in Eq. 1, we obtain a set of functions to fit the experimental D_app_ curves, both at the ensemble average and the single trajectory level. We shall henceforth refer to Table 1 as a reference for the models in questions. In these model fits, we have included a model for free (Brownian) diffusion (Eq. 2), whereby the diffusion is described only by the diffusion coefficient D. Two models for confined diffusion, the approximate version (Eq. 4.1) (52,63) and the exact expression for confined diffusion within permeable square corrals of size L (Eq. 4.2) (50,51) are also included. In these models, the diffusion coefficient is named D_μ_, to indicate that this motion happens within a restricted confinement zone. Notably, the exact formulation contains an infinite sum, which cannot be fit to data analytically. Thus, for this study, we restrict this formula to the case in which k≤39, which converges well to the approximate form (Eq. 4.1) for t>0.5ms. The compartmentalized diffusion models, as it is evident from Eqs. 5.1 and 5.2, are the linear combination of time-independent free (Brownian) motion (Eq. 2) and time-dependent confined diffusion (Eqs. 4.1 and 4.2). The presence of the scaling factor (D_μ_-DM/D_μ_) in Eqs. 5.1 and 5.2 was first introduced in (51) in order to ensure that the intra-compartmental diffusion coefficient (limit of D_app_(t_n_) for t→0) is defined as D_μ_ while the inter-compartmental diffusion coefficient (limit of D_app_(t_n_) for t→∞) is defined as D_M_. We point out that in the analysis adopted, we included the localization uncertainty δ_xy_, as a free parameter in the fitting routine, in order to better estimate for the dynamic nature of this parameter in the detected particle tracks.

**Table 2.** Mathematical expressions for the models considered for the analysis. The following parameters are defined for all diffusion models as follows: MSD is the mean squared displacement, D is the diffusion coefficient, t_n_ is the product of the number of experimental data points, n, and the interval between two consecutive frames, t_lag_, v is the directed flow velocity, L is the average compartment size, D_μ_ is the unhindered, intra-compartmental diffusion coefficient within a confining compartment, D_M_ is the hindered, inter-compartmental diffusion coefficient between confining compartments, *Γ* is a transport coefficient, and α is an the anomaly coefficient where α < 1 for sub-diffusion and α>1 for super-diffusion.

| Diffusion Model | Diffusion coefficient |
| --- | --- |
| | $D(t_n) = \frac{MSD(t_n)}{4 t_n} \quad t_n = n t_{lag}$ |
| 2 – Free (Brownian) Diffusion | $D(t_n) = D$ |
| 3 - Directed motion | $D(t_n) = \frac{v^2 t_n}{4}$ |
| 4.1 - Confined Diffusion (approximate) | $D(t_n) = \frac{L^2}{12 t_n} \left[ 1 - \exp\left(-\frac{12 D_\mu t_n}{L^2}\right) \right]$ |
| 4.2 – Confined Diffusion (exact) | $D(t_n) = \frac{L^2}{12 t_n} \left( 1 - \frac{96}{\pi^4} \sum_{k=1,3,5,\dots}^{\infty} \frac{1}{k^4} \exp\left(-\left(\frac{k \pi}{L}\right)^2 D_\mu t_n\right) \right)$ |
| 5.1 – Compartmentalized (Hop) Diffusion (approximate) | $D(t_n) = D_M + \frac{D_\mu - D_M}{D_\mu} \frac{L^2}{12 t_n} \left[ 1 - \exp\left(-\frac{12 D_\mu t_n}{L^2}\right) \right]$ |
| 5.2 - Compartmentalized (Hop) Diffusion (exact) | $D(t_n) = D_M + \frac{D_\mu - D_M}{D_\mu} \frac{L^2}{12 t_n} \left( 1 - \frac{96}{\pi^4} \sum_{k=1,3,5,\dots}^{\infty} \frac{1}{k^4} \exp\left(-\left(\frac{k \pi}{L}\right)^2 D_\mu t_n\right) \right)$ |
| 6 - Anomalous Diffusion | $D(t_n) = \Gamma t_n^{\alpha-1}$ |
| 7 – Immobile particle | $D(t_n) = 0$ |

Model fits were performed using a weighted nonlinear least-squares fitting routine (NonLinearModelFit[] in Mathematica version 12.0.0.0, Wolfram Research). The D_app_ curves for the model fits were sampled non-linearly, in order to ensure that the points at larger time lags are not overwhelmingly weighted compared to the fewer points at the earliest time lags. This is done as the first few time points are more informative on short-lived events, such as transient confinements. The sampling is thus operated by converting the time axis to a logarithmic scale, and sampling in intervals of length (log_10_ *T* – log_10_ *t*_0_)/(0.5 * *T*/*t*_0_), where T is the maximum time range considered for the analysis. We have chosen to perform our analysis at five different time ranges, that is, five different values of T (5 ms, 10 ms, 25 ms, 50 ms, 75 ms and 100 ms), in order to cover a variety of time regimes with our study.

The most suitable model to describe each particle track is then selected through minimization of the Bayesian Information Criterion (64) for each model whose parameter estimates converge to a non-zero magnitude with a p<0.05 significance level. The condition R^2^>0.9 is taken as fit quality metric, especially at the single trajectory level where the D_app_ curves tend to be most affected by measurement error.

### Compartmentalization metrics

The average confinement time τ_Conf_ (6,65,66), which is a metric that represents the average residence time of a resides inside a compartment, can be defined as:

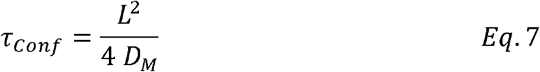

Finally, we include the so-called anomalous diffusion model (Eq. 6), given its widespread adoption in the field. This is a modification to the free diffusion law, whereby the diffusion coefficient has a power-law dependence on time. Accordingly, the multiplication factor is not, in fact, a diffusion coefficient, given the dimensionality mismatch with this quantity, but has rather been called “Transport Coefficient”. In order to quantitatively describe the compartmentalization of the cell membrane, we have adopted another dimensionless quantity, first introduced in (51), called confinement strength, which we indicate as S_Conf_, and is defined as:

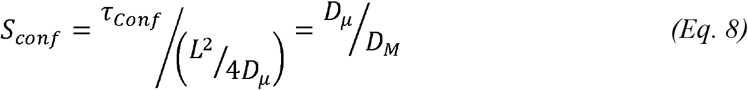

This quantity can be defined as the residence time of a particle, normalized to the time that a freely diffusing particle would spend in the same region, if it also diffuses at the same rate. The two extreme cases for this metric is the case of free Brownian diffusion where D_μ_=D_M_ and S_Conf_=1, and the case of confined diffusion where D_M_=0 and S_Conf_=∞ while the ratio varies continuously in the case of compartmentalized diffusion. Regrettably though, the confinement strength metric cannot readily be used in the case of anomalous diffusion as the limits of t->0 and t->∞ diverge to respectively D_μ_-℞ and D_M_->∞.

### Monte Carlo simulations of 2D diffusion in a heterogeneous lattice

Monte Carlo simulations were performed using custom written routines in MatLab as previously described (8), which can be easily replicated using the Python package described in (61). In brief, in these simulations, we generated fluorescence time traces of 2-dimensional diffusion of single molecules in a heterogeneous corralled environment. The corrals are randomly generated via a Voronoi tessellation algorithm, with randomly selected seeds, to simulate the heterogeneity of the cellular membrane environment. The simulation area was a square with side lengths of 8 to 20□μm (the dimensions of the area are not influential) and the compartmentalisation of this area was implemented as a Voronoi mesh on a uniform random distribution of seed points. We defined the square root of the average compartment area as the average compartment size or length L. The average compartment size (L), defined as the square root of the average compartment area, the hopping probability (Phop) and the free diffusion coefficient (D_S_) completely described our simulation model. Within a compartment the molecules were assumed to diffuse freely while crossing from one compartment to another is regulated by a “hopping probability” P_hop_. This was implemented in the following way: if the diffusion motion (with diffusion coefficient D_S_) would make the lipid to cross the compartment boundary, a random number is generated, and the movement takes place only if this number is above the threshold defined by P_hop_. In all other cases, a new displacement is calculated, where the molecule would be diffusing in the same compartment. In the special case of free Brownian diffusion (i.e. P_hop_ =1) each collision with a compartment boundary results in a molecule crossing to the adjacent compartment. When P_hop_<1, e.g. for P_hop_=1/40, only 1 out of 40 collisions with a compartment boundary results in a molecule crossing to the adjacent compartment. For each condition (D_s_, L_s_, and P_hop_), we simulated N=100 trajectories with 0.5 ms time steps (i.e. a sampling rate of 2 kHz) and a time span of 250 ms (i.e. 500 displacements per trajectory). Subsequently, we added a localization offset δ_rs_ to each localization in the simulated trajectory {x_i_,y_i_}, to simulate the effects of a fixed localization uncertainty in experimental data. These trajectories were subsequently analysed by use of same data analysis pipeline as for the experimental ISCAT data, except that the camera blur correction factor was set to R=0, for obvious reasons.

## Results and Discussion

### Labelling and imaging of cells

In this work, we have used ISCAT microscopy to investigate the lateral diffusion in the apical plasma membrane of live PtK2 cells of artificially incorporated biotinylated phospholipids, DSPE-PEG2000-Biotin, tagged with either Ø20 nm or Ø40 nm streptavidin-coated gold nanoparticles. In order to avoid false detections, whereby a moving particle could be detected, for example, diffusing outside of a cell, the cell membranes were also labelled with a fluorescent lipid (Atto488-DOPE), and simultaneously imaged using the TIRF channel present in our setup (see Materials and Methods). The movies of diffusing particles where then recorded only in the areas where a fluorescent signal corresponding to a cell was detected.

### ISCAT microscopy enables the collection of long, continuous single particle trajectories at 2 kHz frame rates

The main challenge in the data analysis of diffusing particles is the stochastic nature of this phenomenon. Consequently, the robustness in the determination of descriptive physical parameters of any such process by SPT is significantly improved by the availability of long, preferably continuous single particle trajectories. Furthermore, the sampling frequency of the data acquisition needs to be sufficiently rapid to resolve permanent or transient confinements into compartments in the tens to hundreds nanometres range (1,2,11). Using our ISCAT set-up, we have been able to acquire long, continuous, trajectories of diffusing biotinylated lipid analogues (DSPE-PEG(2000)-Biotin), inserted in the plasma membrane of live PtK2 cells, while labelled with either Ø20nm or Ø40nm streptavidin-coated gold particles at a sampling frequency of 2 kHz. This sampling frequency adopted was deemed adequate for detecting transitions in the diffusion mode of the tagged lipids due to the compartmentalization of the plasma membrane, which previous studies at faster frame rates detected in the millisecond time range, with compartment sizes in the hundreds of nanometres (2,3). The ensembles of the track segments are visualized in Fig. 2a for Ø20nm and Fig. 2b for Ø40nm gold nanoparticles. Although this visualization cannot give detailed information on the diffusion mode, or physical quantities thereof, it is still apparent that the gold particle-labelled DSPE lipids on PtK2 cells diffuse from the centre in a broadly symmetrical manner, reflecting the stochastic nature of the long-term motion of the tracked particles.

**Figure 2.**
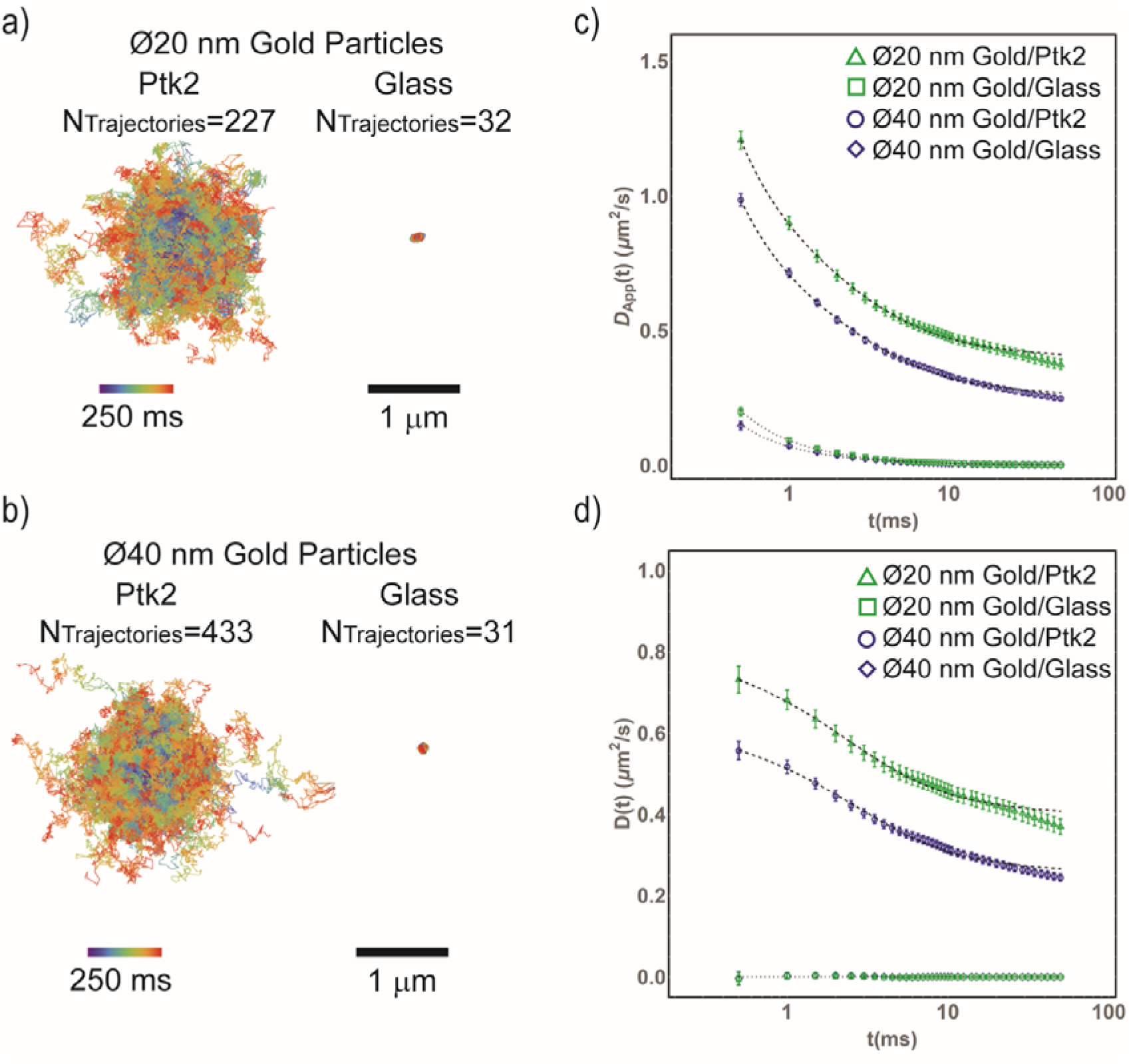
Experimental ISCAT tracking data. a) Superimposed trajectories from time-lapse image acquisitions of diffusing DSPE-PEG(2000)-Biotin lipid analogues in the plasma membrane of Ptk2 cells labelled with Ø20 nm diameter, streptavidin-coated gold nanoparticles, and (right) the same gold nanoparticles immobilized on glass (acquisition frame rate = 2kHz). All trajectories were truncated to 250ms (n=500 displacements) long segments, re-mapped to start at the same position. The color scale indicates the time of each localization (blue to red). b) Same as in a) except that lipid analogues were labelled with the Ø40nm streptavidin-coated gold nanoparticles. c) Comparison between the ensemble average of the D_app_(t_n_) curves obtained from diffusion of Ø20nm gold-tagged lipids on PtK2 cell membranes (green triangles) and Ø40 nm gold-tagged lipids on PtK2 cell membranes (blue circles), and the same size nanoparticles immobilized on glass (Ø20 nm (green squares); Ø40 nm (blue diamonds)). The most likely model of diffusion was the exact compartmentalized diffusion with localization uncertainty (Eq. 5.2) for both nanoparticle sizes, and localization uncertainty (Eq. 7) for Ø20 nm, respectively free diffusion, with a low magnitude diffusion coefficient, with localization uncertainty (Eq. 2) for Ø40 nm gold nanoparticles nanoparticles immobilized on glass (dashed lines). The fit results are shown in Table 3. d) Same data shown in a), but where the localization uncertainty δ_xy_ has been subtracted. Estimation of the localization uncertainty is performed by leaving δ_xy_ as a free parameter when the most likely model is fit to the data in each case (Table 3). The most likely model fits for each curve, corrected for localization uncertainty, are also reported in dashed lines.

### The analysis of the ensemble average D_app_(t_n_) curves are largely invariant for analysis time regimes 0.5ms ≤ t_n_ ≤ T with 10 ms ≤T ≤ 100 ms

Once all the trajectories were collected, we applied the data analysis pipeline to our ISCAT tracking data of Ø20 nm and Ø40 nm streptavidin-coated gold particle-tagged DSPE-PEG2000-biotin lipids on PtK2 cell membranes, and to the same gold particles immobilized on a glass surface. Initially, we decided to analyse the ensemble averages, divided by probe, of the D_app_(t_n_) curves obtained from each trajectory, calculated as in Eq. 1. One of the challenges in the analysis of SPT data is that there is no consensus regarding the amount of points of a MSD curve to be analysed in order to properly describe the diffusion motion of the tracked particle (67). For this reason, we evaluated the dependence on the fit results upon the analysis time range for six different time intervals, 0.5ms ≤ t_n_ ≤ T, where T was set to 5ms, 10ms, 25ms, 50ms, 75ms, or 100ms. At the employed sampling frequency of 2kHz, this corresponds to 10 (2%), 20 (4%), 50 (10%), 100 (20%), 150 (30%), and 200 (40%) points of the 500 localization long segments extracted from the full-length trajectories.

To reiterate, we then fitted the models in Table 1 to the ensemble averages D_app_(t_n_) curves obtained from each kind of sample. From our analysis, we found out that the most likely model to describe the time-dependence of the ensemble averages D_app_(t_n_) curves for both the Ø20 nm and Ø40 nm gold particle-tagged lipids is the compartmentalized diffusion model, in the formulation given by Powles and co-workers (50) (Eq. 5.2), with the simplification that the first 17 terms are considered for the infinite sum (Fig. 2c,d). The resulting fit parameters are reported in Table 3. The second most likely model is the approximate form of the same model (Eq. 5.1), with a significant relative likelihood value. A full summary of the fit parameters for these two models, for every time range, is given in Supplemental Tables S2 and S3. The other models, being much more unlikely according to our selected fit quality metrics, were not included.

**Table 3.** Fit Parameters for the exact Compartmentalized diffusion model fit to the ensemble average D_app_(t_n_) curves for the gold-tagged DSPE biotinylated lipids. The model fit to the data is presented in Table 1, Eq. 5.2, together with an explanation of the fit parameter. The errors reported for the D_μ_, D_M_, L and δ_xy_ refer to the Standard Error of the Fit, as exported from the fitting routine implemented in Mathematica, with the same number of significant digits as the fit parameter. The error for the derived metrics confinement time τ_Conf_ (Eq. 7) and confinement strength S_Conf_ (Eq. 8) is obtained with standard rules of error propagation. The fit parameters here reported refer to model fitting performed on the analysis time range 0.5 ≤ n δ_t_ ≤ 50 ms.

| Sample | $D_\mu$ (±S.E.)<br>($\mu\text{m}^2/\text{s}$) | $D_M$ (±S.E.)<br>($\mu\text{m}^2/\text{s}$) | $L$ (±S.E.)<br>(nm) | $\delta_{xy}$ (±S.E.) (nm) | $\tau_{\text{Conf}}$ (ms) | $S_{\text{Conf}}$ |
| --- | --- | --- | --- | --- | --- | --- |
| Ø20nm Gold<br>nanoparticle-tagged DSPE | 0.87±0.02 | 0.398±0.004 | 110 ±3.0 | 14.2±0.2 | 7.5±0.4 | 2.2±0.0 |
| Ø40nm Gold<br>nanoparticle-tagged DSPE | 0.67±0.02 | 0.256±0.003 | 100 ±2.3 | 13.5±0.2 | 10±0.6 | 2.6±0.1 |

In Fig.2c, we reported the ensemble average of the D_app_ curves for the trajectories collected. It is evident to see how these curves diverge at short time intervals. This is intuitive, given that the expression in Eq 1 has a term with the time lag, t_n_, at the denominator. The fact that the localization uncertainty has an influence on the detected diffusion coefficient of moving particles is well known (62,68). Nevertheless, we have highlighted, by showing the D_app_ curves of immobilized gold nanoparticles, how the localization uncertainty affects all the trajectories, leading to potential overestimation of diffusivity. It is also not sufficient to subtract a constant offset from the curves, as it is usually done in relevant literature (e.g., (2)) since it is demonstrable (62) how the localization uncertainty is variable depending on the diffusion coefficient. Given the uncertainty in estimating, aprioristically or globally, a diffusion coefficient for particles that undergo transient compartmentalization, we have thus opted to introduce the localization uncertainty δ_xy_ as a fit parameter, to have a more precise estimation of its value. Once the model fit has been performed, the δ_xy_ can be subtracted, and the localization uncertainty-corrected D_app_ curves obtained. In Fig.2d, we report the results of this operation. We would like to point out how this operation, applied to the D_app_ curves of the immobilized gold nanoparticles on glass, produces a flat line at 0 μm^2^/s. Naturally, the model that best fits the immobilized gold nanoparticles D_app_ curves is the Immobile particle one (Eq. 7), for which the fit parameters are reported in Supplemental Tables S4 and S5. Finally, we report that the fit-derived δ_xy_ values for the diffusing gold-tagged lipids are consistently larger than those for the immobilized gold nanoparticles (Fig.3d). This should further drive the point that this critical parameter ought not to be established a priori for a single particle tracking experiment, as that will inevitably lead to an underestimation.

**Figure 3.**
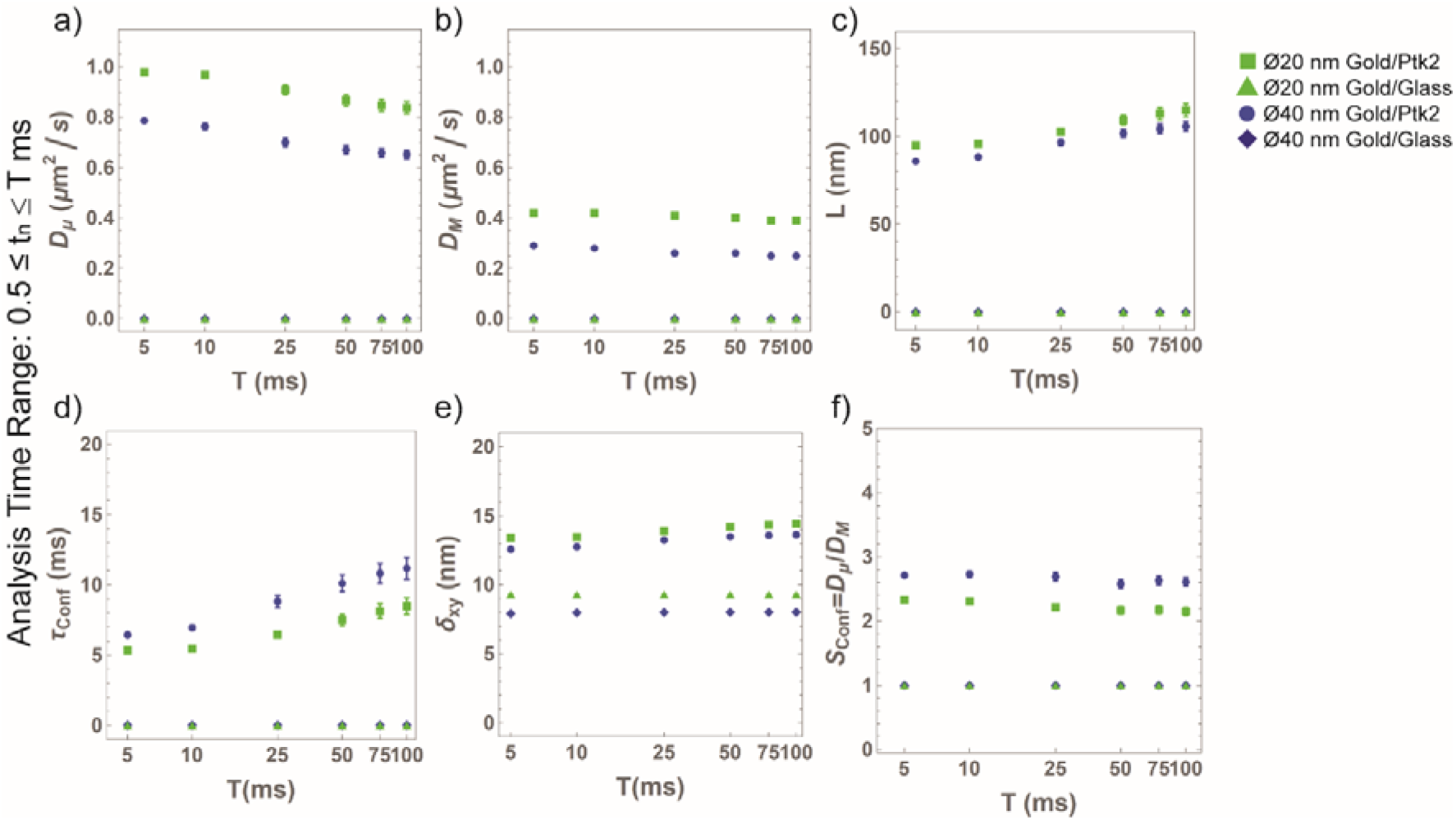
Average Trajectory Analysis Parameters. **Comparison of results emerging from the analysis of the ensemble average D_app_(t_n_) curves**. a–f) The ensemble average D_app_(t_n_) curves obtained from the trajectories of Ø20 nm gold-tagged DSPE lipids on PTK2 cell membranes (black open triangles), Ø40 nm gold-tagged DSPE lipids on PTK2 cell membranes (blue filled triangles), and the same nanoparticles immobilized on glass (Ø20 nm (black filled squares); Ø40 nm (blue open triangles)) were analysed at different time intervals (0.5 ≤ tn ≤ T ms with T= 5, 10, 25, 50, 75, and 100 ms). We plot here the resulting fit parameters, obtained from the fitting the most likely diffusion model to the data, and the resulting confinement strength metrics. The values shown here shown here are: a) the unhindered diffusion coefficient at time t=0, D_μ_, b) Timeindependent (“free”) diffusion coefficient D_M_, c) the confinement size L, d) the confinement time τ_Conf_=L^2^/(4 D_M_), e) the localization uncertainty δ_xy_, and the confinement strength S_Conf_= D_μ_ /D_M_.

From Fig. 3, it is possible to appreciate that the fit parameters are somewhat stable for analysis time ranges longer than 10 ms. Shorter analysis time ranges cannot fully capture the compartmentalization dynamics of the target gold-tagged DSPE lipids. Thus, we report (Table 3) only the values of the fit parameters for the ensemble averages obtained through the compartmentalized diffusion models at the analysis time range 0.5ms ≤ t_n_ < 50ms to offer a representation of the compartmentalization dynamics observed on the PtK2 cell membrane. This corresponds to approximately 20% of the total data points in each trajectory segment considered by our analysis, putting this choice below the “rule of thumb”, which prescribes not to use more than 25% of the total track duration (67,69), and the approach adopted by Kusumi and co-workers, using only the first few points for the analysis (70,71).

### The magnitude of the diffusive motion parameters, but not the motion type, are dependent on the probe size

The fit parameters here reported paint an interesting picture of the dynamics of the gold-tagged DSPE lipids on the PtK2 cell membrane. First of all, we notice that the difference in size produces a generalized reduction in the diffusivity of the target lipids, both in the D_M_ and D_μ_ parameters. Nevertheless, it is remarkable that the model describing the diffusion is still the same, which would suggest that the introduction of a large tag only slows down the particle dynamic, without introducing artefacts in the detected diffusion motion. This conclusion is consistent with our previous findings (36), which show a similar effect in model membranes (i.e. Supported Lipid Bilayers) and live cells (6–8).

The detected compartment sizes L differ slightly between the two probe sizes (L=100±2.3nm for the Ø40nm gold nanoparticle-tagged lipids, and L=110±3.5nm for the Ø20nm gold nanoparticle-tagged ones), and, most notably, differ by a factor larger than two from reported results on a similar system (2,3). One reason from this can be attributed to the fundamental difference in the data analysis strategy here adopted, and specifically, the handling of the localization uncertainty. In the present study, this quantity is handled as a fit parameter. In fact, a finite localization uncertainty can lead to very strong artefacts on the apparent diffusion coefficient D_app_(t_n_), especially when short frame times are involved, as it can be deduced from Eq. 1. The localization uncertainties estimated this way cannot be compared to those reported in similar studies, due to the difference in the estimation methods (see Supplemental Note 1), nevertheless they fall close to accepted values from comparable observations, that is, δ_r_ = 13.5±0.2 nm and δ_r_ = 14.2±0.2 nm for the Ø40nm and Ø20nm gold-tagged DSPE respectively.

In Table 3, we also included the confinement time τ_Conf_ (Eq. 7), a quantity also known as average residency time within a compartment. Being directly related to the L and D_M_ parameters (Eq. 7), this quantity obviously differs between the two samples here considered, and would agree with the observation that generally the diffusion rate of the lipids tagged with the larger Ø40nm gold nanoparticle experience is overall slower, as can be intuitively understood. However, we find that a more fitting quantity to describe the compartmentalization of the plasma membrane is represented by the confinement strength (Eq. 8), first introduced in (51), and here reported as SConf. Interestingly, the calculated values of this metric are quite close between the Ø20 nm and Ø40 nm gold-tagged DSPE (2.2±0.1 and 2.6±0.2, respectively). This, again, would indicate that while the size of the gold nanoparticle probe could have an influence on the detected motion of the target DSPE lipid, but it doesn’t fundamentally alter the lateral diffusion dynamics of lipids in the plasma membrane. This conclusion is consistent with our previous findings (36), and is compatible with the observation that the viscosity of the water-based medium into which the gold nanoparticle extends offers less restriction to diffusion than the high-viscosity, obstacle-rich cellular membrane environment (72).

### The single trajectory analysis reveals the full heterogeneity of the lipid motion on the plasma membrane of Ptk2 cells

The analysis of the ensemble average D_app_, exposed in the previous section, provides an efficient and immediate way to evaluate the overall most likely diffusion model for all the trajectories detected. Most landmark SPT studies on similar samples are restricted to such data analysis, often with much smaller sample sizes than the present work (e.g., (1,2)). However, SPT, as an analysis method, has the potential to reveal the extent of diffusion heterogeneity down to the single trajectory level. We thus applied our analysis pipeline to each single molecule track. In order to streamline the analysis, we took the decision of only adopting the “exact” models from Table 2, where the decision is possible. The distributions of the fit parameters results are visualized in Figure 4, while the summary statistics are reported in Table 4. From Fig.4a-b, it is possible to see that the relative fractions of the most representative models stabilize for time intervals 0.5ms ≤ t_n_≤ T with 25 ≤ T ≤ 100 ms.

**Figure 4.**
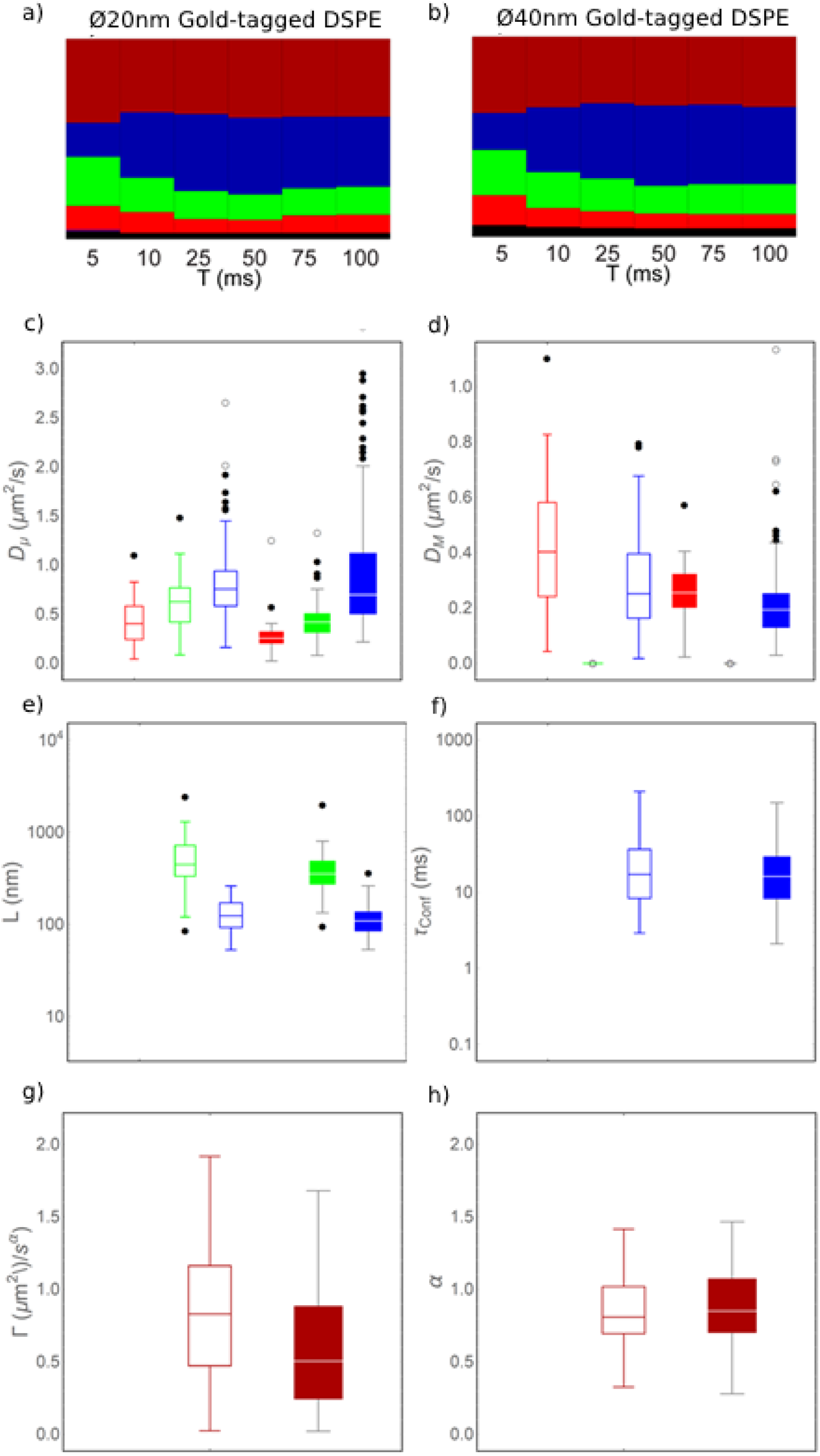
Model fitting of the Apparent Diffusion Coefficient curves for each single trajectory. a-b) Visual representation of the relative fractions of the ensemble of trajectories divided by most likely diffusion model at different analysis time ranges (T, with the notation for time range 0.5 ≤ t_n_ ≤ T), for DSPE lipids labelled with Ø20 nm gold nanoparticles (a) and Ø40 nm gold nanoparticles (b). The models are: anomalous diffusion (Eq. 6, dark red), exact compartmentalized diffusion (Eq. 5.2, blue), exact confined diffusion (Eq. 4.2, green), free diffusion (Eq. 2, bright red). In black, the fraction of trajectories for which no model offers a good plot under the chosen fit quality metrics. c) Distributions of the fit parameter D_μ_ obtained by fitting the most likely model for each trajectory. The empty bars correspond to the Ø20 nm gold-tagged DSPE lipids, while the full bars to the Ø40 nm gold-tagged DSPE lipids. The models included are exact compartmentalized diffusion (Eq. 5.2, blue), exact confined diffusion (Eq. 4.2, green), free diffusion (Eq. 2, bright red). The anomalous diffusion model is not included, since it doesn’t incorporate a diffusion coefficient in its formulation. The free diffusion model is included to allow a comparison between all models containing diffusion coefficients. d) Distributions of the fit parameter D_M_, obtained as in c). The models represented are only exact compartmentalized diffusion (Eq. 5.2, blue) and free diffusion (Eq. 2, bright red). e) Distributions of the fit parameter L, obtained as in c). The models represented are only exact compartmentalized diffusion (Eq. 5.2, blue) and exact confined diffusion (Eq. 4.2, green), in which this parameter appears. f) Distributions of the τ_Conf_ metric calculated for the trajectories whose most likely model is the exact compartmentalized diffusion (Eq. 5.2), divided by probe (empty bars for Ø20 nm gold-tagged DSPE lipids, full bars for the Ø40 nm gold-tagged DSPE lipids). g) Distributions of the fit parameter Γ for the ensembles of single trajectories best described by the anomalous diffusion model (Eq. 6). The trajectories are divided by the probe employed (empty bars for Ø20 nm gold-tagged DSPE lipids, full bars for the Ø40 nm gold-tagged DSPE lipids). h) Distributions of the fit parameter αfor the ensembles of single trajectories best described by the anomalous diffusion model (Eq. 6). The trajectories are divided by the probe employed (empty bars for Ø20 nm gold-tagged DSPE lipids, full bars for the Ø40 nm gold-tagged DSPE lipids).

**Table 4.**
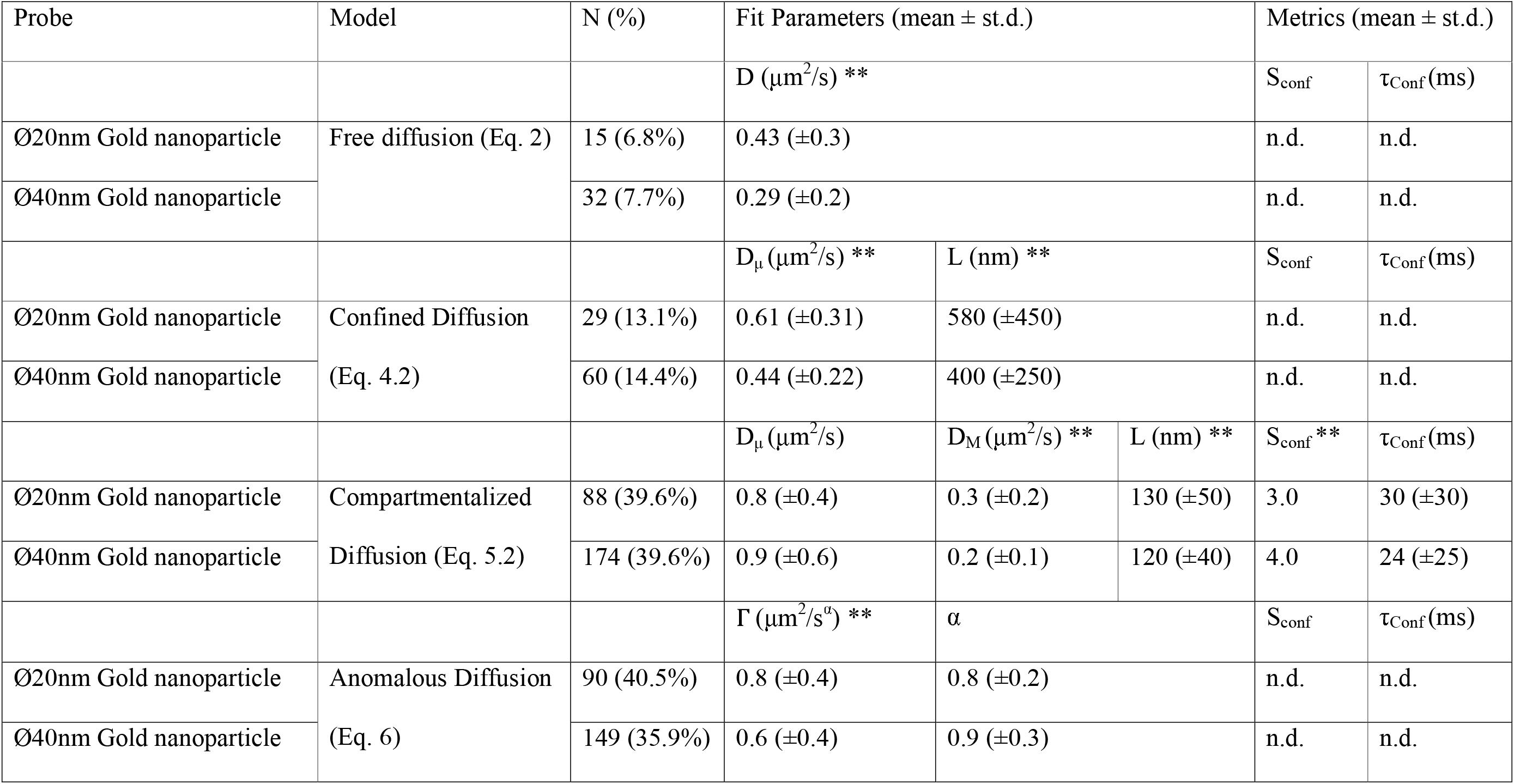
Summary statistics of the fit parameters of the single trajectory analysis for the two species of probes targeting biotinylated DSPE Lipids. The diffusion models mentioned in the “Model” column were fit to the D_app_ curves derived from each single 500 localization segment derived from the original particle trajectories, divided by probe size, for the gold nanoparticle-tagged DSPE lipids. In the column N (%), we report the number of single particle trajectories, and the fraction of the total number of 500-localization segments that it represents (Table 1). The number of fit parameters in the eponymous column is, of course, variable, and reflects the equation mentioned in the Model column. We indicate with the ** notation, the fit parameters whose distribution is different according to the Kolmogorov-Smirnov test, with a level of significance <0.05. The fit parameters here reported refer to model fitting performed on the analysis time range 0.5 ≤ n δ_t_ ≤ 50 ms.

Our analysis showed that the relative fractions of the most likely lateral motion model of the single trajectories, for both species of nanoparticles, were roughly constant in the time analysis ranges of 0.5ms ≤ t_n_ ≤ T for 10 ≤ T ≤ 100 ms (Fig. 4a-b). The exact confined diffusion model appears more dominant at the shortest time window (0.5ms ≤ t_n_ ≤ 5ms), as this short analysis time interval may not be sufficient to fully capture the transition from intra-compartmental diffusion to inter-compartmental diffusion. Therefore, we further elaborate on this single particle trajectory analysis for the results obtained using as time range 0.5ms ≤ t_n_ ≤ 50ms, for consistency with the analysis of the ensemble average D_app_(t_n_) curves. We report the exact fractions of trajectories best described by each model in Table 4. To fully appreciate the differences between these distributions, we have compared the parameters across samples through the Kolmogorov-Smirnov (KS) test, but comparing only among populations of the same models.

First of all, we compare the magnitude of the diffusion coefficients D_μ_ and D_M_ across the different models, and samples (Fig. 4c-d). For the trajectories classified via the free diffusion model, it is very easy to appreciate how the values for the diffusion coefficient D are significantly different, with the Ø20 nm gold nanoparticles-tagged DSPE diffusing faster (D = 0.4±0.3μm^2^/s) than the Ø40 nm gold nanoparticles-tagged DSPE (D = 0.3±0.2μm^2^/s). The significance of the difference carries over to the values for D_μ_ for both the confined and compartmentalized diffusion models, but not to the D_M_ for the compartmentalized diffusion model (D_M_ = 0.3±0.2μm^2^/s and D_M_ = 0.2±0.1μm^2^/s for the Ø20 nm and Ø40 nm gold nanoparticles-tagged DSPE, respectively).

The distributions of the parameter L (Fig. 4e) suggest that there may be two distinct levels of compartmentalization in the plasma membrane of live cells. One level of compartmentalization is highlighted by the fraction of trajectory classified as Compartmentalized diffusion (Eq. 5.2). In this case, we can see that L=130±50 nm for the Ø20 nm gold nanoparticles-tagged DSPE, and L=120±40 nm for the Ø40 nm gold nanoparticles-tagged DSPE. In fact, the particles best described as purely confined (Eq. 4.2), experience confinements on quite large spatial scales, specifically L=580±450 nm for the Ø20 nm gold nanoparticles, and L=400±250 nm for Ø40 nm gold nanoparticles. The lipid trajectories best described by this model exhibit a macroscopic diffusion coefficient D_M_ = 0, thus representing a fraction of particles that eventually becomes immobilized when observed for 250ms. The distributions of these values are significantly different (p<0.05), under the KS test (Table 4).

A very interesting result emerges when observing the distribution of the compartmentalization metrics τ_Conf_ and S_Conf_ which, again, can only be calculated in the case of compartmentalized diffusion. While the distributions of τ_Conf_ (Fig. 4f) across samples is not significantly different, the distributions of S_conf_, instead, is (Table 4).

We find that the anomalous diffusion model (Eq. 5), best describes an almost constant portion, around 40%, of the single particle trajectories. Given that this fraction is seemingly unaffected by the analysis time interval, might indicate a certain lack of sensitivity of this model for the exact diffusion behaviour of the target molecule. While the coefficients of transport Γ are very significantly across the two probe species (Fig. 4g), we report that this difference does not carry over the anomaly coefficient α (Fig.4h). In fact, the difference between the relative distributions of the Γ parameter is significant (P>0.05), under the KS test, while those of α are not.

This single trajectory analysis has revealed that, underneath what is revealed by the the analysis of the ensemble average D_app_(t_n_) curves, there is a wealth of complexity that should not be overlooked. The study of the ensemble averages offers undoubtable advantages, such as a less noisy signal, or the convenience of quickly summarizing the collective behaviour of a sample. Nevertheless, the ensemble average diffusivity should, at least, be performed on a sizeable sample size, in order to include as much of the heterogeneity observed herein as possible (72).

### Validation of data analysis by comparison to simulated diffusion on a heterogenous lattice

Apparent compartmentalization and trappings can appear in diffusion, as a result of the natural variability in random walks. This has been known since the first observation of single particle diffusion (73,74). More recently, it has been suggested that Monte Carlo simulations of diffusions should be used as control for the experimental data (74,75). Therefore, we compared the results from our experiments to simulated trajectories on a compartmentalized surface as described in the Materials and Methods section, and in (6,8,76). This procedure enabled us to directly compare our observations with our hypothesis for the mechanisms generating the heterogeneities of diffusion on the plasma membrane, namely its compartmentalization. In order for the simulations to be as realistic as possible, we generated trajectories of particles diffusing on a randomly generated environments with semi permeable corrals designed by a Voronoi tessellation algorithm, with an indicative compartment cross section L as control parameter.

We simulated 100 tracks on such environments, with different sets of input parameters (see Materials and Methods), chosen to match as closely as possible the ensemble average D_app_ curves of the experimental data (data not shown). The experimental ensemble average ISCAT data for Ø20 nm gold tagged DSPE could be well approximated by simulated data with the following parameters: P_Hop_=0.06, D_S_=1.0 μm^2^/s, L_S_=120 nm, and δ_r S_=16 nm. On the other hand, the data for Ø40 nm gold tagged DSPE could be very well approximated by simulations with parameters P_Hop_=0.04, D_S_= 0.8 μm^2^/s, L_S_=120 nm, and δ_r S_=16 nm. We then analysed the ensemble average D_app_ curves obtained from the simulated trajectories with the same data analysis pipeline as the ensemble average of experimental data. Unsurprisingly, the simulated trajectories were also well described by the compartmentalized diffusion models (Eq. 5.2). We report in Table 5, the fit parameters related to the 0.5 ms ≤ T ≤ 50 ms time interval, with the fit parameter of the corresponding experimental data in the same time range. The fit parameters for the other time ranges are reported, for completeness, in graphical form in Fig. 5, and in Supplemental Tables S6 and S7. The closeness between the model fit parameters between the simulations and the experimental data is quite striking, especially considering that the generation of the simulated trajectories only stems from a very elementary description of the environment, i.e., its division in somewhat regularly sized, but not shaped, compartments. In particular, the compartmentalization metrics τ_Conf_ and S_conf_ appear to be very closely matched, which highlights how well the compartmentalization model matches the sample as probed by our experiments. However, we must point out how the P_hop_ and the D_S_ differ between the simulated datasets. This might be due to the larger Ø40 nm gold nanoparticles slowing down the detected DSPE lipid diffusion dynamics, and in doing so, altering also the probability of the particle to change compartments.

**Figure 5.**
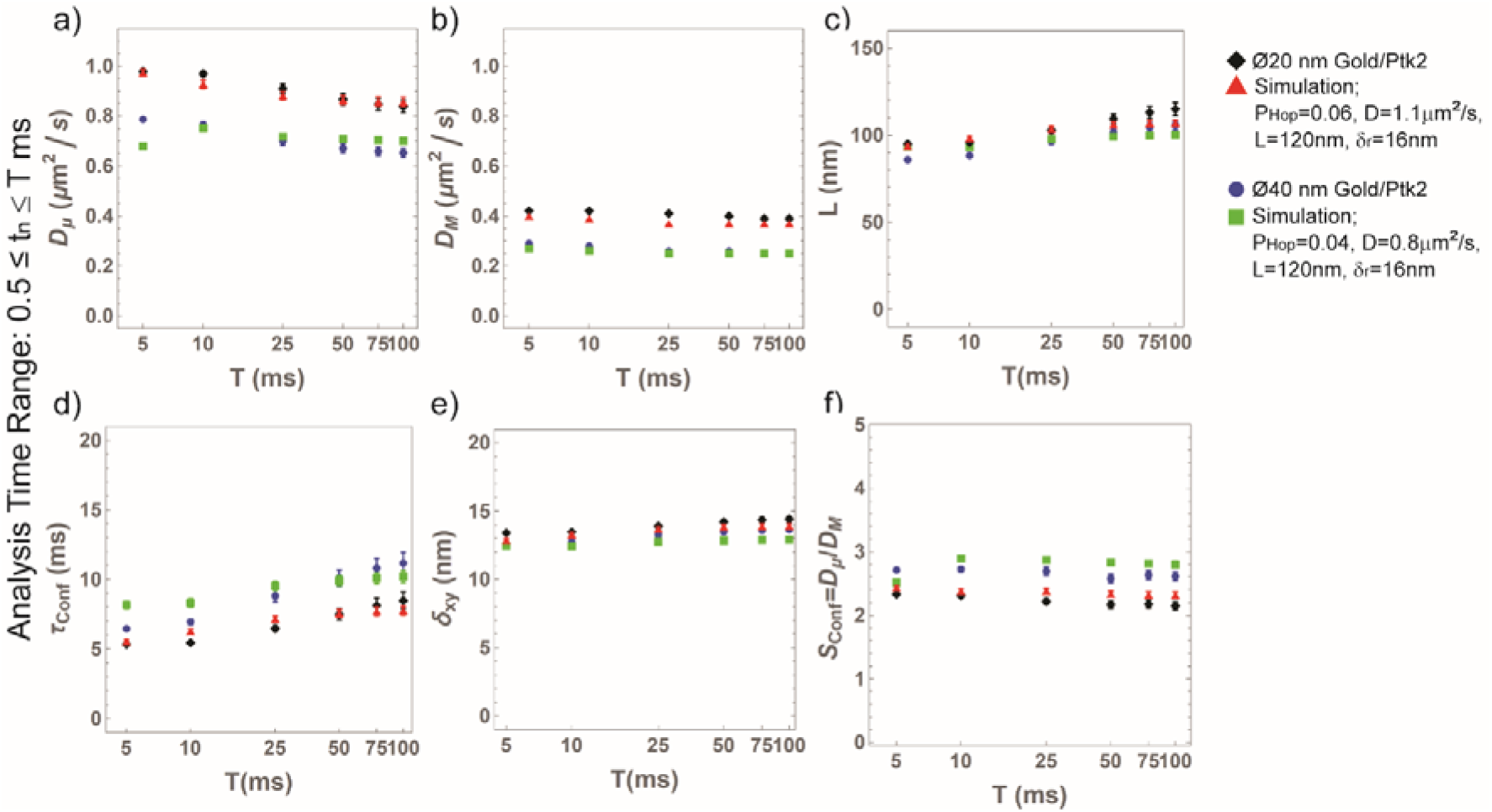
Comparison between average fit parameters between experimental data and matching simulated data. a–f) The ensemble average D_app_(t_n_) curves obtained from the trajectories of Ø20 nm gold-tagged DSPE lipids on PTK2 cell membranes (black diamonds), simulated trajectory data for compartmentalized diffusion in a heterogenous lattice with simulated parameters P_Hop_=0.06, D=1.1 μm^2^/s, L=120 nm, and δ_r_=16 nm (red triangles), Ø40 nm gold-tagged DSPE lipids on PTK2 cell membranes (black circles), and simulated trajectory data with simulated parameters P_Hop_=0.04, D=0.8 μm^2^/s, L=120 nm, and δ_r_=16 nm (green squares) were analysed at different time intervals (0.5 ≤ t_n_ ≤ T ms with T= 5, 10, 25, 50, 75, and 100 ms). The most likely model of diffusion for all data sets and time intervals was the exact compartmentalized diffusion (Eq. 5.2) with localization uncertainty. We plot here the resulting fit parameters, obtained from the fitting the most likely diffusion model to the data, and the resulting confinement strength metrics. The values shown here shown here are: a) the unhindered diffusion coefficient at time t=0, D_μ_, b) Time-independent (“free”) diffusion coefficient D_M_, c) the confinement size L, d) the confinement time τ_Conf_=L^2^/(4 D_M_), e) the localization uncertainty δxy, and the confinement strength S_Conf_ = D /D_M_.

**Table 5.** Comparison between experimental data and matching simulations. Exact Compartmentalized Diffusion model (Eq. 5.2) fit parameters of the ensemble average Apparent Diffusion Coefficient curves of the experimental data, and the fit parameters obtained from the corresponding matching simulations. Model fitting is performed on the analysis time range 0.5 ≤ n δ_t_ ≤ 50 ms. ^†^Values of uncertainty <0.005, reported with two significant digits for consistency.

| Sample | $D_{\mu}$ (±S.E.) | $D_M$ (±S.E.) | L (±S.E.) | $\delta_r$ (±S.E.) (nm) | $\tau_{\text{Conf}}$ (ms) | $S_{\text{Conf}}$ |
| --- | --- | --- | --- | --- | --- | --- |

**Table 5. Comparison between experimental data and matching simulations.**
| | ($\mu\text{m}^2/\text{s}$) | ($\mu\text{m}^2/\text{s}$) | (nm) | | | |
| --- | --- | --- | --- | --- | --- | --- |
| Ø20nm Gold<br>nanoparticle-tagged DSPE | $0.87 \pm 0.02$ | $0.40 \pm 0.00^\dagger$ | $110 \pm 3.0$ | $14.2 \pm 0.2$ | $7.5 \pm 0.4$ | $2.2 \pm 0.1$ |
| $P_{\text{hop}} = 0.06$, $D_S = 1.1$<br>$\mu\text{m}^2/\text{s}$, $L_S = 120\text{nm}$, $\delta_{rS} =$<br>16 nm | $0.86 \pm 0.02$ | $0.37 \pm 0.00^\dagger$ | $106 \pm 2.0$ | $13.8 \pm 0.2$ | $7.6 \pm 0.3$ | $2.3 \pm 0.1$ |
| Ø40nm Gold<br>nanoparticle-tagged DSPE | $0.67 \pm 0.02$ | $0.26 \pm 0.00^\dagger$ | $100 \pm 2.3$ | $13.5 \pm 0.2$ | $10 \pm 0.6$ | $2.6 \pm 0.1$ |
| $P_{\text{hop}} = 0.04$, $D_S = 0.8$<br>$\mu\text{m}^2/\text{s}$, $L_S = 120\text{nm}$, $\delta_{rS} =$<br>16 nm | $0.71 \pm 0.01$ | $0.25 \pm 0.00^\dagger$ | $99 \pm 1.0$ | $12.9 \pm 0.1$ | $9.9 \pm 0.4$ | $2.9 \pm 0.1$ |

The simulation framework laid out in this work, proves useful also in appreciating the effect of environmental parameters on the detected diffusion. In Fig. 6a-b, we show what the application of a constant offset δ_s_ to each localization, as a substitute for the localization uncertainty δ_xy_, has on the ensemble average D_app_ curves. It is evident that, when the offset δ_s_ is larger, then the D_app_ diverges at short t_n_. These simulations were performed using the same frame time as the experimental data as interval between localizations, but it is easy to extend this reasoning to the case of faster framerates. This effect is fortunately easy to correct (Fig. 6b): when the diffusion models are fit to the simulated D_app_ curves, and a δ_xy_ value is obtained as for the experimental data, this can then be subtracted to reliably obtain the true D(t_n_) curves. Given that the localization uncertainty increases for moving particles (62,68), when the localization uncertainty is estimated from immobilized probes (e.g., (1,2)), this will inevitably lead to some degree of overestimation of the diffusion coefficient.

**Figure 6.**
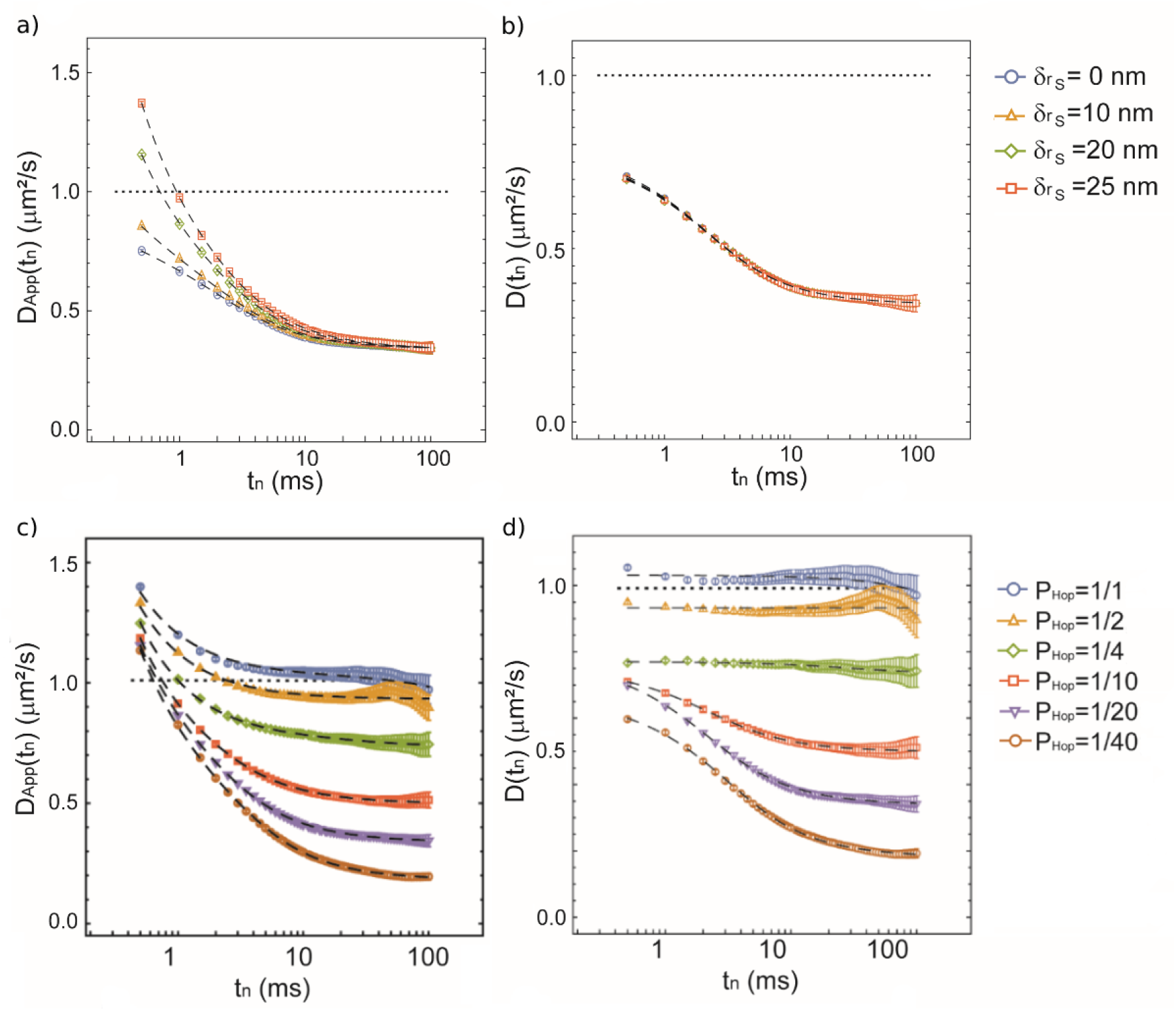
Effect of the localization offset δ_rS_ and P_hop_ on the simulated diffusion of particles. a) Ensemble average D_app_(t_n_) curves of sets of 100 simulated trajectories where a constant offset δ_rS_ (see legend) has been added to each localization before calculating the D_app_. The trajectories were simulated as described in the Materials and Methods, with the rest of the simulation parameters being: P_Hop_=1/20, D_S_ = 1.0 μm^2^/s and L_S_ = 120 nm. b) Same data as in a), after subtraction with a localization uncertainty δxy obtained from fitting the ensemble average curves with model for compartmentalized diffusion (Eq. 5.1) to the D_app_(t_n_) curves in a). c) Ensemble average D_app_(t_n_) curves of sets of 100 simulated trajectories, with different values of P_hop_ (see legend). The trajectories were simulated as described in the Materials and Methods, with the rest of the simulation parameters being: δ_rS_=20 nm, D_S_ = 1.0 μm^2^/s and L_S_ = 120 nm. d) Same data as in c), after subtraction with a localization uncertainty δ_xy_ obtained from fitting the ensemble average curves with model for compartmentalized diffusion (Eq. 5.1) to the D_app_(t_n_) curves in c).

We also evaluated the effect of different P_hop_ on the D_app_(t_n_) curves detected (Fig. 6cd). In this instance, we can see how weak compartmentalization (P_hop_>1/4) is essentially indistinguishable in behaviour from free diffusion (Phop=1), only resulting in a net decrease in the observed diffusivity compared to the a priori established D_S_. On the other hand, the compartmentalization dynamics start becoming more apparent for lower values of P_hop_, which present a more noticeable transition between the two diffusion regimes, macroscopic and microscopic. Once again, we must point out how striking the difference is between the original (Fig. 6c) and the localization uncertainty-corrected datasets (Fig. 6d).

Finally, another observation can be extracted from the simulation data. In fact, the datasets thus originate also present trajectories which are best described by different diffusion modes, apart from the compartmentalized one. This is to be expected, given the stochastic nature of the simulated trajectories (72,77). However, we observed how the L parameters originating from these trajectories are much larger than those obtained from the trajectories best described by compartmentalized diffusion models (unpublished data). This might serve to explain the same differences obtained from the single particle trajectories, which evidently originate from particles that are, by accident, “trapped” during the observation time. This only reinforces the concept that single particle diffusion data ought to be carefully analysed at multiple levels.

### The confinement strength metric enables direct comparison of cell membrane diffusion across different techniques

We have thus far adopted the S_Conf_ metric (Eq. 8) as a tool to compare the observed compartmentalization dynamics across gold nanoparticle size, and in relation with the Monte Carlo simulations used to confirm our observation. However, it is also possible to use this parameter to draw the connection between simulated data and the experimental data not only from this work, but also from studies of lipid diffusion in related literature (6–8). In our simulation framework, this parameter is strongly connected, but not equivalent, to the “hopping probability” P_Hop_, and it should provide with a representative descriptor of the physical landscape where the tracked particles are diffusing. By calculating this metric for the ensemble average data presented in this work, we obtain the values S_conf_=2.2± 0.1 for the lipids tagged with the Ø20 nm gold nanoparticles, and S_conf_=2.6 ± 0.1 for the lipids tagged with the Ø40 nm gold nanoparticles (Table 5). From these results, we can confirm our observation that the larger probe size has a stronger influence on the detected motion of the target particle, resulting in a higher measured confinement strength. Nevertheless, when we compare these values with the S_conf_ obtained from the analysis of the ensemble average D_App_(t_n_) data from the matching simulated trajectories (Table 5), we find a close resemblance with the corresponding values obtained from the experimental data.

In the broader context of comparison of the present data with relevant literature, the S_conf_ parameter also shows its potential to classify the diffusion motion of target lipids on cellular membranes. In particular, we found that it is possible to quite effectively extend this framework to other methods to detect diffusion dynamics, such as Fluorescence Correlation Spectroscopy (FCS), and its combination with super-resolution STED microscopy (STED-FCS). However, while in the present case the values of D_μ_ and D_M_ are readily available from the compartmentalized model fits such is not the case for techniques such as (STED-)FCS. Thus, different definitions must be found. For the experiments reported in (7,8), where diffusion is detected via STED-FCS, we set the equivalent of the parameter D_μ_ as the diffusion coefficient detected via STED-FCS with the smallest detection spot (i.e., with the highest depletion laser power), whereas the D_M_ would be the diffusion coefficient detected using the conventional diffraction limited spot. A special case is represented by the experiment in (6), in which the diffusion of a fluorescent lipid analogue (Atto647N-DPPE), is detected by conventional FCS on the surface of Ptk2 cells, in the presence and absence of CK666, which inhibits Arp2/3 mediated actin crosslinking, and thus the compartmentalization of the plasma membrane. In this case, we consider the D_M_ is the diffusion coefficient measured on the cells treated with the drug, where the diffusion should be unrestricted, whereas the D_μ_ is the same parameter measured in the absence of the drug, and thus in a compartmentalized environment.

In Figure 6, we plotted the values for S_conf_ obtained from the experiment thus far presented and from the aforementioned related studies against the values of P_Hop_ estimated by matching simulations. In fact, in the present study and in (6–8), the experimental data was matched to the same kind of simulations described in this study. Thus, while it is not possible to give a direct estimation of the parameter P_hop_ from the experiments, it is possible to give a close estimate, that can be related to the S_conf_. To these points, we overlaid the values of S_conf_ derived from the the analysis of the ensemble average D_app_(t_n_) curves of simulated trajectories with different values of P_Hop_ (other simulation parameters: D_S_=1.0 μm^2^/s, L_S_=120 nm, and δ_r S_=20 nm) (Fig. 7). Corresponding relevant parameters are reported, for completeness, in Table 6.

**Figure 7.**
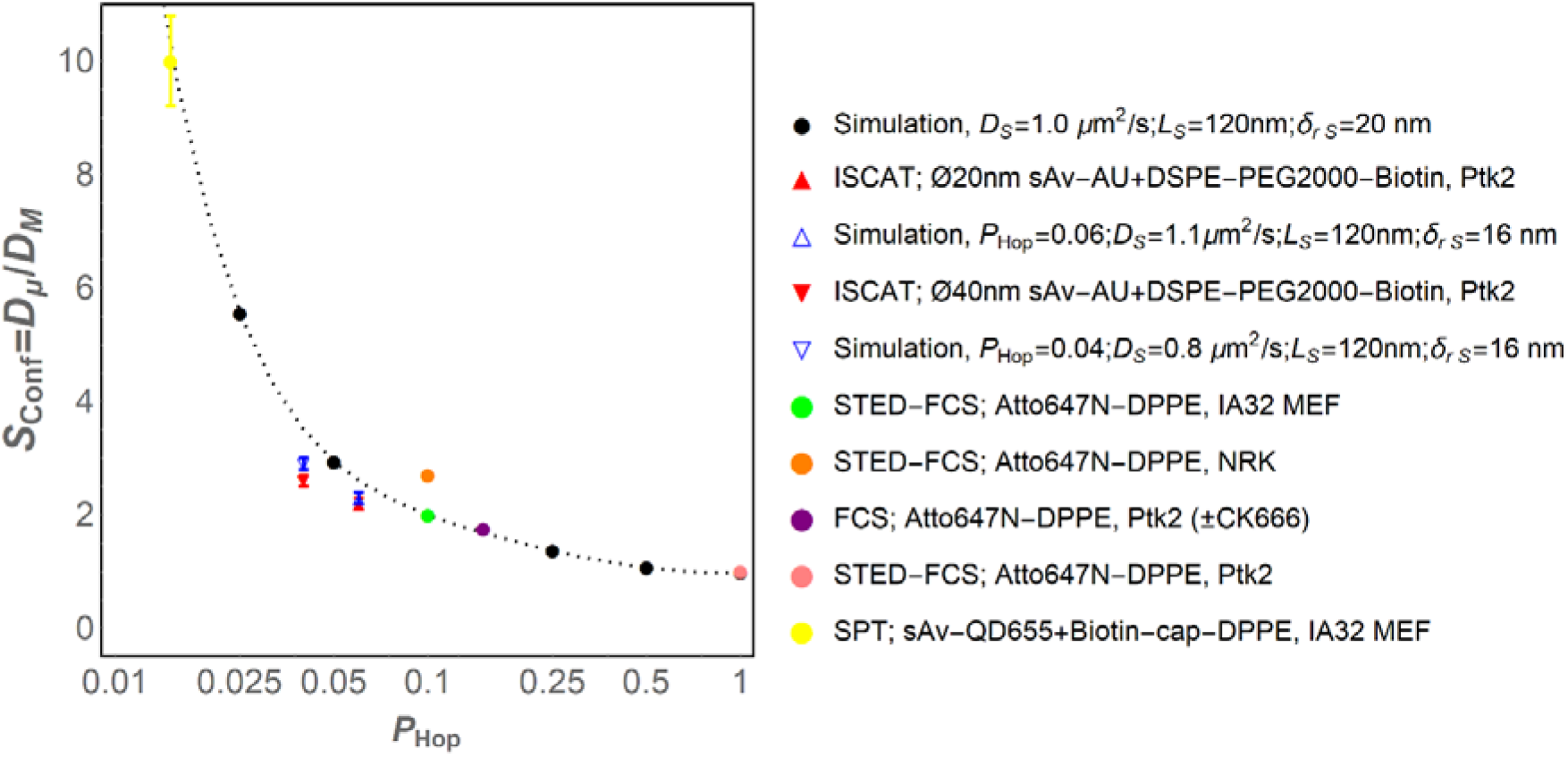
Comparison of ISCAT experimental results to our previous lateral diffusion in plasma membrane of live cells by STED-FCS, FCS, and SPT. To put the results from this study into context with previous related studies, we have plotted the confinement strength S_Conf_=D_μ_/D_M_, as determined by comparative analysis of simulated trajectory data for compartmentalized diffusion, versus the hopping probability, P_Hop_, for data as indicated in the legend. All data is also reported in Table 6.

**Table 6.**
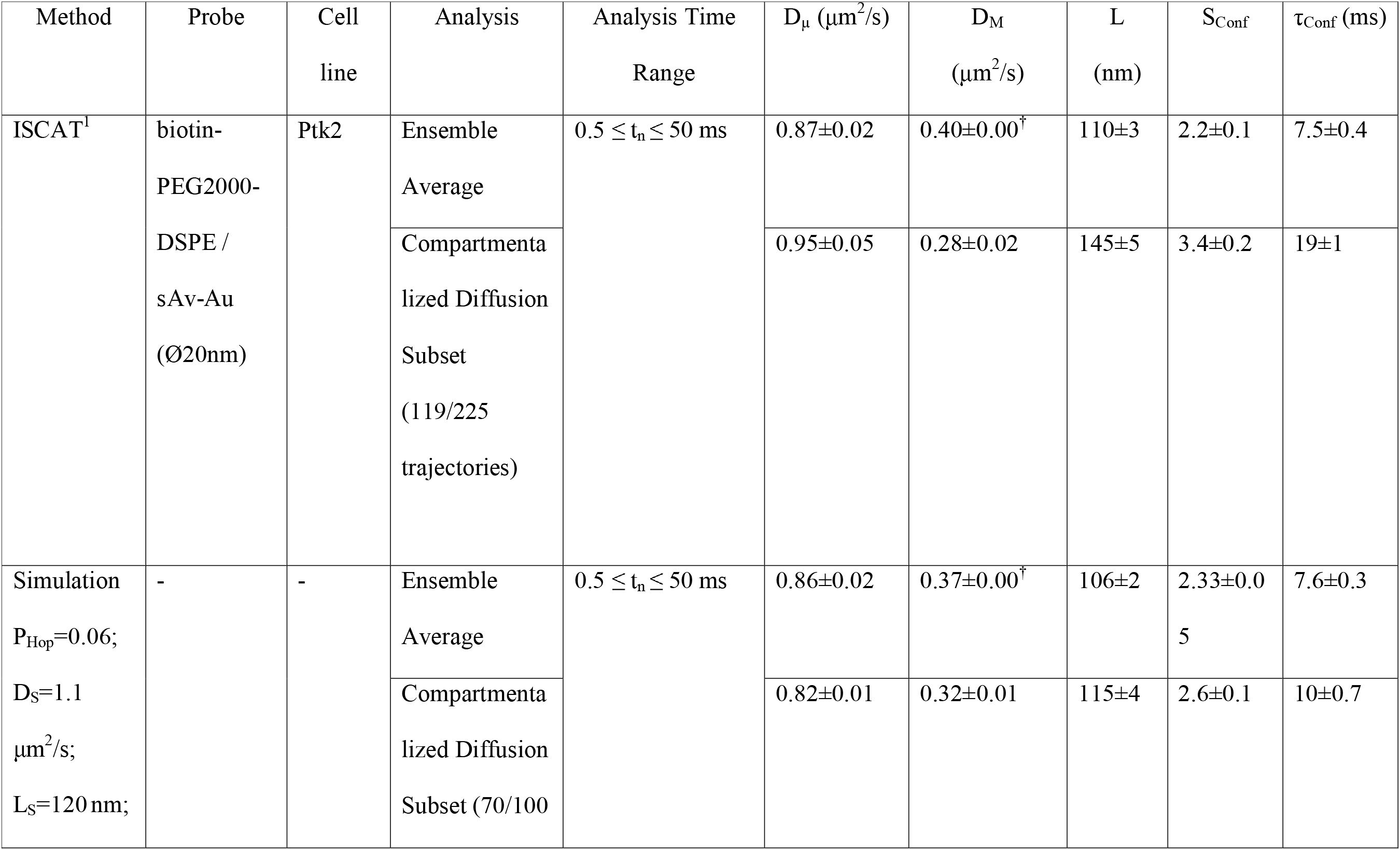
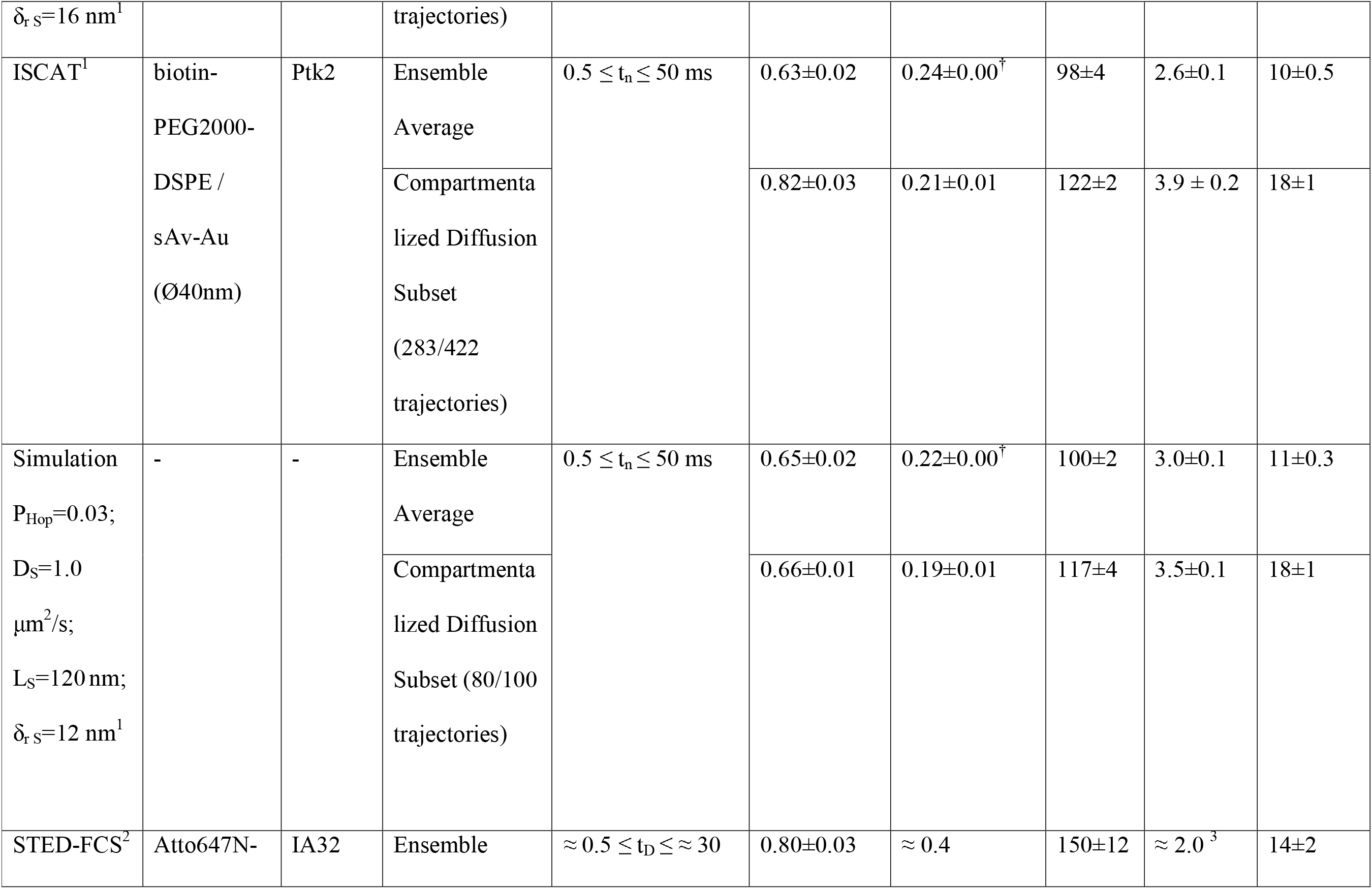
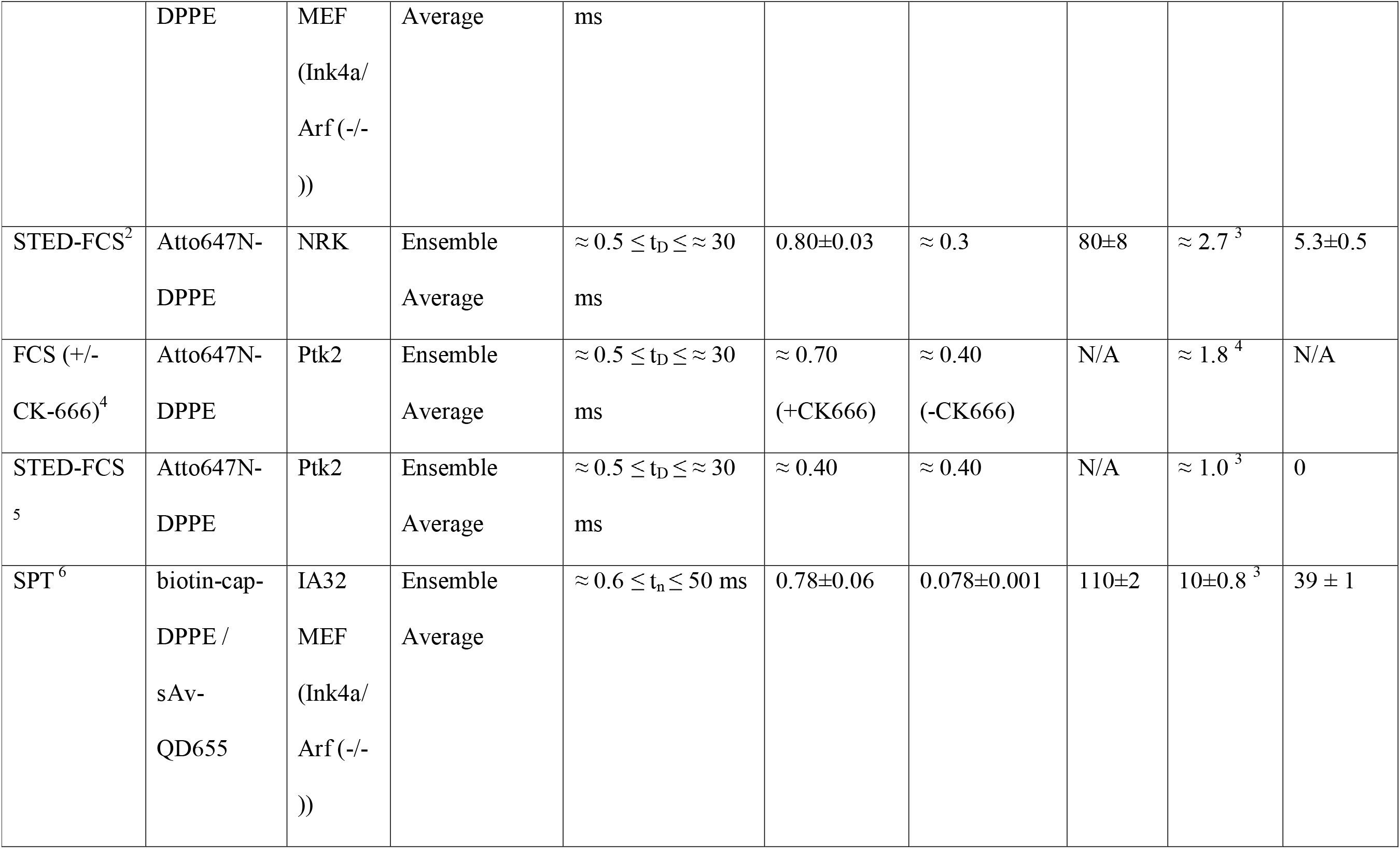
Comparison of values for Diffusion coefficients, S_conf_, τ_Conf_ and other relevant parameters obtained from the diffusion experiments and simulations from this study, with those obtained from related literature.^1^. Data featured in this work.^2^ Data featured in (8).^3^ For STED-FCS measurements, we consider D_μ_ the diffusion coefficient calculated from data acquired at the highest available STED power (corresponding to an approximate lateral resolution of 50 nm for the specific studies), whereas we have defined D_M_ as the diffusion coefficient calculated from data acquired at confocal lateral resolution (~250 nm). Thus, the confinement strength, S_Conf_, for STED-FCS is defined as S_Conf_ =D_μ_/D_M_=D_STED_/D_Confocal_.^4^ For conventional FCS measurements with and without Arp2/3 specific inhibitor CK-666, we have defined D_μ_ as the diffusion coefficient calculated from measurements in the presence of 100□μM CK-666, whereas D_M_ is the extracted diffusion coefficient for data acquired in the absence of CK-666. Thus, the confinement strength, S_Conf_, for FCS in this case is defined as 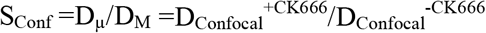.^5^ Data featured in (7). ^6^ Data featured in (6). ^†^Values of uncertainty <0.005, reported with two significant digits for consistency.

The values of S_conf_ thus obtained can be used to relate amongst each other measurements originating from vastly different experiments, both in terms of technique and probe used. For example, it is possible to compare a mostly freely diffusing probe, such as Atto 647N-DPPE on Ptk2 cells measured (6) (S_conf_=1), with a mostly confined probe, such as Biotin-cap-DPPE tagged with streptavidin-coated quantum dots on the membrane of IA32 MEF cells (6) (S_conf_=10±0.8), despite the first one being measured with STED-FCS, and the latter through fluoresce-based SPT. This metric is more effective than the confinement time τ_Conf_, since the ratio between the microscopic and macroscopic diffusion coefficients (Eq. 8) manages to normalize over the influence of the probe, which might reduce the diffusivity of the target molecule through phenomena unrelated to the lateral diffusion.

This metric might in the future prove useful when comparing measurements across techniques on the same sample, or when comparing how different lipids feel the influence of membrane compartmentalization in a different way, due to their properties (e.g., degree of saturation, etc.)

## Conclusions

In this work, we have adopted ISCAT microscopy as a method to probe the compartmentalization of the living cell membrane, through Single Particle Tracking studies of gold naπoparticle-tagged, biotinylated lipid analogues (DSPE). We defined and significantly expanded a data analysis pipeline framework for analysis of single particle trajectories that has previously been presented in (6). One of the main purposes of the present work was to present a clear data analysis methodology, grounded in theory and mindful of localization uncertainty artefacts, which could be implemented to analyse similar dataset. The data analysis methodology we presented here is based on the observation of the apparent diffusion coefficient D_app_(t_n_), a quantity derived directly from the raw mean squared displacements (MSD) (Eq. 1), after suitably correcting it for motion blur (68) and environmental oscillations. With this revised analysis protocol, we are also able to correct the raw apparent diffusion coefficient D_app_(t_n_) from the effect of the localization uncertainty, which we estimated as a fit parameter in our data analysis routine. The importance of such correction has been highlighted in Fig. 6.

From the formulation of the diffusive motion models adopted (Eq. 2-7), we have been able to directly extract the physical properties underlying the plasma membrane environment, effectively using the tracked particles as a probe. In particular, from the results of the data analysis on the ensemble averaged D_app_(t_n_) data, the compartmentalized diffusion model, as defined by (50,51), emerges as the most descriptive of the models considered. This model (Eq. 5.2) is representative of an environment where the diffusing particles are divided into compartments surrounded by a partially permeable barrier. Thus, the movement of the particles can be separated into a short-range, time-decaying component, typical of the faster motion inside the compartments, and a long-range, time-invariant component that is representative of the slower motion between compartments. The analysis of the ensemble average D_app_(t_n_) of the trajectories also revealed that the larger Ø40 nm gold nanoparticles, while noticeably slowing down the lipid dynamics observed, do not alter the diffusion mode significantly in the case considered. We also analysed each of the single trajectories, taken singularly, following the same protocol. This revealed a high degree of heterogeneity in the diffusion modes detected, and the physical parameters extracted. Nevertheless, the adopted protocol is also completely viable for single particle trajectory, thus revealing an analysis method that could prove very informative, as it can directly address relevant physical parameters relative to the structure of the plasma membrane.

Another strategy we employed to further our understanding of the plasma membrane environment is to compare the experimental data to simulated trajectories. To this end, we employed a simulation framework in which a two-dimensional space is corralled into compartments of fixed average size L_S_ and random shape, over which Brownian particles diffuse at a fixed rate D_S_ and have a probability of “hopping” from one compartment to the other P_Hop_. This is effectively a direct implementation of the “hop” diffusion model described analytically in (50). With an appropriate choice of initial parameters, we thus obtained sets of trajectories, that we analysed with the same analysis pipeline as the experimental data. The fit parameters obtained from the D_app_(t_n_) data of these simulated datasets closely match those obtained from the experiments, and so do the resulting fit parameters. We thus have crossvalidated the conclusions made from the experimental data, and confirmed the model of plasma membrane herein considered.

We should note that the results for lipid diffusions on PTK2 cell membranes here reported are in stark contrast with similar observations made in the past (1,2), albeit at faster frame rates (40-50kHz), and using a piece-wise analysis strategy, where the first points of the MSD curve where analysed separately from the rest. In particular, in the work of Murase et al. (2), for similar experimental conditions, the compartment size was estimated at 44nm, with an average residence time of 1.5ms. The origin of this differences can be attributed to, first of all, the differences in the estimation of the localization uncertainty. This is because it directly affects the estimation of the apparent diffusion coefficient of the particle. Secondly, the estimation of these quantities is made through very different means, and thus a direct comparison between the quantities is very challenging.

The simulation framework also allows us to incorporate this work in a broader context, involving more studies and data from comparable experiments (6–8). For this, we extended the definition of confinement strength (S_conf_), originally proposed in (51) to be compatible with FCS and STED-FCS. Using this parameter, we were able to correlate the results obtained with other similar diffusivity measurements in different cell lines, conditions and techniques, to sets of simulated trajectories specifically generated. A clear pattern emerged, where the S_conf_ of the experimental data approaches the S_conf_ obtained from matching simulations, where the P_hop_ is chosen for best matching the D_app_(t_n_) of simulated and experimental trajectories. While this result may seem obvious, we would like to stress that this result emerges from considering the compartmentalization as the only source of diffusion heterogeneity for an otherwise Brownian-diffusing particle. Finally, although the procedure should be refined in order to attain wider applicability, it retains the potential to be a defining feature for future plasma membrane diffusivity experiments, being technique-agnostic and providing an interesting descriptor for the physical characteristics of the cell membrane environment.

The results here described confirm that the semi-permeable compartmentalization of the cellular membrane is largely responsible for the deviation of the observed diffusion from pure Brownian behaviour. However, additional phenomena might contribute to these observations, such as membrane topology (12,13,78), the presence of membrane proteins such as CD44 (14), and naturally, the presence of lipid nanodomains. Clearly, additional observations would be needed to discern between these contributions effectively.

## Acknowledgements

The authors greatly acknowledge the Engineering and Physical Sciences Research Council (EPSRC) and Medical Research Council (MRC) for supporting the DPhil project of FR within the Oxford-Nottingham Biomedical Imaging Centre of Doctoral Training (ONBI-CDT) (Grant No. EP/L016052/1). Further, we acknowledge support by the EPA Cephalosporin Fund (Bio-ISCAT project), the MRC (Grant No. MC_UU_12010/unit programs G0902418 and MC_UU_12025), the Wellcome Trust (Grant No. 104924/14/Z/14 and Strategic Award 091911 (Micron)), MRC/BBSRC/EPSRC (Grant No. MR/K01577X/1), the Wolfson Foundation (for initial funding of the Wolfson Imaging Centre Oxford), the John Fell Fund, and the Deutsche Forschungsgemeinschaft (DFG, German Research Foundation; under Germany’s Excellence Strategy – EXC 2051 – Project-ID 390713860, and project number 316213987 – SFB 1278 (project C05)), as well as the Jena Center for Soft Matter (JCSM). The authors would like to acknowledge the support of Dr. Erdinc Sezgin and Dr. Dilip Shrestha for their scientific advice, and the Kukura Group (Oxford University) for the technical support in the initial phases of microscope development.

## Author Contributions

C.E. and B.C.L. planned the research work. F.R. and B.C.L. designed the experiments. F.R. conducted the experiments, designed and constructed the microscopy setup. B.C.L. and F.R. performed the data analysis. F.R., B.C.L. and C.E. wrote the article.

## Competing Interests

The authors declare no competing financial interest.

## Notes

### Competing Interest Statement

The authors have declared no competing interest.

### Summary of Updates

A major revision compared to the previous version. Enhanced for readability, clarity and overall more concise. New models for diffusion have been included, and the definition of the confinement strength metric has been made consistent with previous literature.

