## Supplemental Material for "Quantification of live cell membrane compartmentalization with high-speed single lipid tracking through interferometric Scattering Microscopy"

**Supplemental Note 1: ISCAT microscopy enables robust particle detection at high SNR on live cells**

Particle detection in ISCAT microscopy is hindered by the presence of spurious reflections in the optical setup (1–3). We therefore applied a temporal median filter to the final image. Briefly, a stack of one hundred images is collected while the sample is moved by the user. If the operation is performed correctly, i.e., the sample is displaced enough while still being in focus, the median filter obtained from this image stack therefore will contain only the fixed contribution to the signal (4). By using this image as background signal, it is possible to dramatically improve the contrast of the sample in the final detected time-lapses. However, the aforementioned median filtering operation is seldom not sufficient to obtain enough particle contrast to ascertain the position of the particle. This is especially true in the case of samples with high scattering backgrounds such as cells. For this reason, we applied a second background reduction strategy, where we subtract the average intensity projection of the detected, median-filtered time lapse. In this way, the high-scattering cell landscape is mostly filtered out, facilitating the detection of the target gold nanoparticles (Supplementary Figure S2).

To allow comparison with previously published results, we estimated the localization performance of the setup using multiple methods (Supplemental Table S1.1). First of all, as described in (5–7), the measurement was carried out by measuring the FWHM of the distribution of relative distances between two immobilized gold nanoparticles on glass (also see Materials and Methods). This yielded a localization precision of 2.6 nm, which is in line with previously reported results (5,7,8). In our opinion, however, this may represent an underestimation, since the particles are localized singularly, and their displacements are not calculated relative to an integral reference point. This, in turn, may lead to an overestimation of the diffusion coefficient at short timescales. Another method we employed, similar to that presented in (9,10), involves determining the standard deviation of the coordinates of localizations for immobilized gold nanoparticles. In this case, we obtained values of 12.9±2.5nm and 11.3±2.3nm for Ø20nm and Ø40nm gold nanoparticles respectively, once again comparable to previously reported measurements (9,11,12).

| Method for calculation | Data Set | Most Likely Model | δ_x_ (nm) | δ_y_ (nm) | δ_xy_ (nm) |
| --- | --- | --- | --- | --- | --- |
| Mean (±s.t.d.) of Standard Deviation of Localizations (Fig.2a & b) | Ø20nm gold nanoparticles on Glass (N=32) |  | 11.2±2.8 | 6.4±1.3 | 12.9±2.5 |
|  | Ø40nm gold nanoparticles on Glass (N=31) |  | 9.2±1.7 | 6.5±3.1 | 11.3±2.3 |
| δ_xy_ (±s.e) from model fitting the most likely model to the ensemble averages D_app_(t_n_) curves trajectory analysis for Most Likely Model (Fig.2) | Ø20nm gold nanoparticles on Glass (N=32) | Confined diffusion (Eq. 4.2, Table 2) | n.d. | n.d. | 9.3±0.1 |
|  | Ø40nm gold nanoparticles on Glass (N=31) | Confined diffusion (Eq. 4.2, Table 2) | n.d. | n.d. | 8.1±0.1 |
| δ_xy_ (±s.e) from model fitting the most likely model to the ensemble averages D_app_(t_n_) curves trajectory analysis for Most Likely Model (Fig.2)  (Table 3) | Ø20nm Gold nanoparticle-tagged DSPE lipids on PtK2 cells (N=229) | Compartmentalized diffusion (Eq. 5.2, Table 2) | n.d. | n.d. | 14.2±0.2 |
|  | Ø40nm Gold nanoparticle-tagged DSPE lipids on PtK2 cells (N=433) | Compartmentalized diffusion (Eq. 5.2, Table 2) | n.d. | n.d. | 13.5±0.2 |

**Table S1.1. Localization uncertainty estimates as determined by different approaches.** The last two rows of the table above, refer to the value from the localization uncertainty as estimated by the data analysis methodology proposed in this text, whereby the localization uncertainty δ_xy_ is derived from the model fitting operation. In this Table, for brevity, we report uniquely the values for δ_xy_ for the obtained through model fitting at the 0.5 ≤ n δt ≤ 50 ms analysis time range. Further values at different time ranges can be found in Supplemental Tables S2 to S4.

**Supplemental Note 2: Analysis of Single Particle Tracking Data**

**2.1 Analysis of Single Particle Trajectories**

In order to thoroughly analyse and extract correct information from the collected data, we have refined the data analysis protocol previously introduced in (13,14). The distinguishing features of this analysis protocol is the treatment of the localization uncertainty and the implementation of a robust, data-driven statistical method of single trajectory classification based on the most likely model of diffusive motion from a set thereof. Furthermore, in order to form a robust analysis pipeline with constant statistical sampling of measured displacements, we have restricted our analysis to single trajectories that contained at least 500 consecutive localizations, thus corresponding to a trajectory duration of 250 ms at our sampling rate of 2kHz. Furthermore, we truncated all trajectories into segments of 500 consecutive localizations.

We calculated the Mean Squared Displacements (MSD), for each single trajectory from

$$MSD (t_{n})=\frac{1}{N-n}\sum_{i=1}^{N-n} {(r_{i+n}-r_{i})}^{2} (Eq. S1)$$

where r_i_ is the particle position at time t_i_, for all available displacements at a given time lag interval t_n_=n δt, where δt is the interval between two successive observations of the same particle, in this case 0.5ms, resulting from a sampling frequency, 1/ δt, of 2kHz. In the text, we refer to “single trajectory analysis” when the data arising from each single trajectory is analysed, and to “ensemble average” analysis when the MSD values of the trajectories belonging to the same dataset are averaged and then analysed.

The conventional data analysis strategy is to subsequently fit the experimentally determined MSD(t_n_) dependence for a particular analysis time range, t_start_ ≤ t_n_ ≤t_stop_ to a relevant model of diffusive motion such as the expression Brownian (free) diffusion on a two dimensional plane, i.e.

$$\mathrm{MSD}\left( t_{n} \right)=4 D t_{n} \left( 1-\frac{2R}{n} \right)+ 4 {\delta_{xy}}^{2} (Eq. S2)$$

where we have also included two factors that are necessary for experimental MSD data. These correction factors are: 1) an additional correction factor, (1-2R/n), to account for motion blur as a consequence of particle motion during the camera integration time (15), and 2) a constant term, 4 δ_xy_^2^, to account for the dynamic localization uncertainty by which each particle position can be determined. The motion blur correction term, which primarily affects the first few data points n, depends on the mode of illumination of the sample. In the case of full frame averaging, with exposure time t_exp_ = 0.227ms, as employed here, R=1/6 (t_exp_/ δt) = 1/6 (0.227 ms/ 0.5 ms) (12,15,16). The localization error term, ${\delta_{xy}}^{2}$ which contributes a constant y-offset, primarily depends on the signal-to-noise of the microscope set-up where ${\delta_{xy}}^{2}$is the dynamic localization uncertainty. These correction factors are an absolute requirement for fast sampling rates as these factors make the trajectories highly non-linear even when simple free diffusion is involved (15,16).

While Eq. S2 is suitable for quantitative analysis, we find it much more informative, due of the complexities that are introduced by the above correction terms, to instead transform the MSD(t_n_) data to the same units of [length^2^/time] as the diffusion coefficient for our case of 2D diffusive motion on a plane as.

$$D_{app}\left( t_{n} \right)=\frac{MSD(t_{n})}{4 t_{n}\left( 1-\frac{2R}{n} \right)} (Eq. S3)$$

The right-hand side of Eq. S3 can be thought of as an apparent diffusion coefficient, D_app_(t_n_) from which it is now possible to directly evaluate the time dependence of the diffusive process from the trajectory data even in the absence of curve fitting to any specific model.

The data, transformed through Eq. S3, was subsequently analysed by least squares non-linear curve fitting, weighed to a range of theoretical models for diffusive motion, all of which directly incorporated the camera blur corrected effect of localization noise as shown in Eq. 2. The generic relationship for the data models that we used for this analysis is obtained by substituting Eq. S2 in Eq. S3:

$$D_{app}\left( t_{n} \right)=D\left( t_{n} \right)+\frac{{\delta_{xy}}^{2}}{t_{n}\left( 1-\frac{2R}{n} \right)} (Eq. S4)$$

where we have split the calculated apparent diffusion coefficient D_app_(t_n_) into its two distinct contributions: 1) the diffusive process D(t_n_), and 2) a camera blur-corrected localization error term, in which δ_r_ is the localization uncertainty.

The diffusion component D(t_n_) in Eq. S4 takes on different formulations to describe a range of theoretical diffusive motion models. In this work we have considered a total of six plausible models for the diffusion component, which can be found in Table 2 in the main text, to which we shall refer in the rest of the present discussion.

The free, or Brownian, diffusion model (Eq. 2), manifests when no obstacle to the diffusion is felt by the particles. We consider two separate models for confined diffusion, respectively the approximate (Eq. 4.1) and exact (Eq.4.2) forms for diffusion in a compartmentalized environment (17–19). The compartmentalized diffusion models, in its approximate (Eq.5.1) and exact forms (Eq 5.2) (17–19) can be thought of as the sum of the aforementioned models. These models present the obvious advantage to being directly dependent of physical parameters relevant for the description of diffusion in a compartmentalized environment. The observed diffusion coefficient within the confining compartment is D_μ_, while the diffusion coefficient between compartment is here defined as D_M_. The parameter L, that appears both in the confined and compartmentalized diffusion models, refers to the average compartment size.

We included the so-called anomalous diffusion model (Eq. 5), given its widespread usage in the community. This expression aims to account for all heterogeneities in diffusion using a power law, and classifying the motion through the exponent α. In particular, for α>1, the motion is classified as “superdiffusion”, for α<1 “subdiffusion”, and “diffusion” for α=1. In this expression, the factor Γ is usually called “transport coefficient”, given its dimensionality mismatch with a diffusion coefficient. The adoption of such model, while it offers a fast avenue for evaluation of diffusion data in heterogeneous environment, does not however offer a direct connection with relevant physical quantities, unlike other models considered. Finally, the immobile particle model (Eq. 6) was included, as a way of filtering particles erroneously classified as diffusing. Initially, our analysis included also a directed motion model. However, none of the trajectories, in any of the time ranges ever exhibited that behaviour, according to our analysis, and is therefore left out of the discussion for simplicity.

These expressions for D(t_n_) could then be substituted in Eq. S4 to obtain models that account for the dynamic localization uncertainty. Using Eq. S3, we could obtain values for D_app_(t_n_), to which model fitting can be performed. The least-squares fitting routine was implemented in Mathematica (version 12.0.0.0; Wolfram Research) using the NonlinearModelFit[] command, weighing inversely to the variance of the D_app_(t_n_), with a Confidence Level of 0.99.

In this analysis, we have further directly included the dynamic localization uncertainty, δ_xy_, as a free parameter in the least squares fit. As a consequence, it may appear as though the localization uncertainty of the measurements is higher, and thus the quality of the experiment worse, than previously recorded experiments. However, this is a different approach to the majority of the model fitting routines presented in relevant literature, where this quantity is either not taken into account during the derivation of parameters relevant to describe the diffusive motion of the particles, or just used as a baseline value to subtract to the entire dataset (20). This is a crucial point, as it is evident from Eq. S4 that the localization uncertainty, especially for kHz sampling rates and above, could produce an apparent diffusion coefficient that is much higher than the “true” one, when not properly taken into account. Another issue is represented by the estimation of the localization error. In other cases the localization error is estimated by measuring the degree of variability in the relative distance of two immobilized tags (7). In the case of ISCAT microscopy, the majority of studies (e.g. (5,21)) report the localization uncertainty of the setup as the average of the standard deviations of repeated localization of immobilized particles (or molecules), determined by fitting the intensity profiles of detected particles to a 2D spatial Gaussian (Supplemental Note 1). The advantage of estimating the localization uncertainty from the diffusion data is that a more realistic estimation is offered, and that the effect of this source of measurement error is fully accounted for in the analysis.

**2.2 Statistical evaluation of the most likely diffusion model**

In this work, we have chosen to adopt a data-driven diffusion motion classification that does neither require *a priori* knowledge nor assumptions on which diffusive motion model to assign to a particular data set. This approach is a refined extension to our previous work (13), with refinements aimed at increasing the robustness of the approach in order to extend the applicability to the analysis of single trajectories. The challenge of analysing single trajectory data in this context stems from the fact that single trajectories are in general much noisier than the ensemble average of each dataset’s trajectories, as the averaging step involves significantly fewer displacements.

In this extended approach, we determined the most likely diffusion model, from the set of plausible models (Eqs. 2-7), by calculating the Bayesian Information Criterion (BIC) from the results for each data model, according to the formula (22):

$$BIC\left( model \right)=n\ln\left( \frac{RSS}{n} \right)+k\ln\left( n \right) (Eq. S5)$$

where n is the number of data points used for the fitting, RSS is the residual sum of squares, and k is the number of free parameters in the fit. The first term in Eq. S5 is a measure of the Goodness of Fit while the second term adds a penalty function that scales linearly with the addition of each free parameter to the model. Thus, while normally a model with a larger k would inevitably result in a fit with a smaller RSS, the use of BIC as a classification metric allows to evaluate the models more equitably. A smaller BIC value equates to better fit quality. A more intelligible metric of how well each model fits to the experimental data is the relative likelihood, which can be derived from the BIC by using the formula:

$Relative Likelihood=Exp\left[ \frac{{BIC\left( model \right)}_{Min}-BIC\left( model \right)}{2} \right] (Eq. S6)$

where BIC(model)_Min_ is the smallest BIC value from the model fits. By definition, the model whose fit minimizes the BIC has a Relative Likelihood of 1 while all other models have Relative Likelihood <1, making it the most descriptive diffusion model from the array of plausible models defined above.

To enhance the robustness of this approach, we have introduced a second goodness-of-fit metric. We ensure that all free parameters in a given model fit converged to non-zero value with a significance level of p<0.05. The t-statistic values are determined as the fit parameter estimates divided by the standard errors, and the p-values reported are the two‐sided p‐values for the t-statistic, with the null hypothesis being that the parameter is equal to zero. This is introduced given that the parameters considered, especially the Diffusion coefficients, tend to assume very low values. Using these two metrics, the most likely model to describe each trajectory therefore has to satisfy two main conditions: minimizing the BIC (i.e., the highest Relative Likelihood), and that the fit parameters are significantly different than zero. If the most likely fit does not satisfy the second condition, the model with the second-smallest BIC, but which satisfies the t-test condition is then selected, and so on. Finally, if no model is able to simultaneously satisfy both conditions, then the trajectory is excluded from further analysis. Additionally, in the case of single trajectory analysis, which naturally suffers from greater noise than the ensemble average of all the curves, we also require that the coefficient of variation (R^2^) for each diffusion model fit has to satisfy R^2^>0.9 for the trajectory to be included in the analysis. This modified data analysis approach establishes a robust unbiased statistical methodology for determining the most likely type of diffusive motion model, from a particular given set of evaluated motion models, however our approach cannot provide judgement on non-tested models.

**2.3 Sampling of D_app_ points in analysis pipelines**

One of the challenges in the analysis of SPT data is that there is currently no consensus with regards to the amount of data points which should be analysed in order to describe the nature of the diffusion motion of the tracked particle (16,23,24). Therefore, we have performed curve fitting across six different time intervals defined as 0.5 ms ≤ t_n_≤ T, with T equal to 5ms, 10ms, 25ms, 50ms, 75ms, and 100ms. For T=5ms and 10ms, all the values of D_app_(t_n_) (10 and 20, respectively) were used in the fitting operation. In the remaining cases, we have instead sampled the data logarithmically. To do this, we first converted the time values to a natural logarithmic scale, and resampled the time points at fixed intervals of width$\left( \log_{10} T-\log_{10} t_{0} \right)/\left( 0.5*T/t_{0} \right)$. For example, for T=50ms (corresponding to n=100 data points), we selected time points that were spaced (Log[50ms]-Log[0.5ms])/75 apart. After rounding to the closest values and removal of duplicate time points, this results in 46 roughly evenly spaced time points on a log scale where n={1, 2, 3, 4, 5, 6, 7, 8, 9, 10, 11, 12, 13, 14, 15, 16, 17, 18, 19, 20, 22, 23, 24, 26, 28, 29, 31, 33, 35, 37, 40, 42, 45, 48, 51, 54, 58, 61, 65, 69, 74, 78, 83, 88, 94, 100} that we used for the fitting. This effectively serves to place extra emphasis on the initial time points, where we observe the greatest change in magnitude of the apparent diffusion coefficient, due to the influence of the localization error and due to the transient confinements of the particle tracjectories.

**Supplemental Note 3: Validation of data analysis pipeline by comparison to simulated trajectories for diffusion in a heterogenous lattice**

To validate our experimental data analysis approach and results, we also analysed simulated trajectories for diffusion in a heterogeneous lattice, generated through the Voronoi tessellation algorithm, with a characteristic average compartment size, L_S_, as previously described (25). These simulations, although simple, are an extension of the infinite array of square corrals model considered in (17), which serves as the theoretical foundation for the models of compartmentalized and confined diffusion (Table 2) we are considering for the model fits. The irregularities and randomness introduced by the Voronoi algorithm should serve to represent the same effects, found in actual plasma membranes.

In these simulations, molecules are diffusing on a surface with a diffusion coefficient D_S_*.* The surface is divided into compartments, and the particles can go from one to the other with a certain “hopping probability” P_hop_. The parameter P_hop_ can be thought as directly linked to the “permeability” ℘ from (17), where P_hop_=1 would correspond to ℘→∞, whereas P_hop_=0 to ℘=0. We find the parameter P_hop_ more practical to describe our results, and we have thus adopted it in this study. To better approximate experimental conditions, we further add a constant offset δ_r S_ to the trajectory positions, in place of the localization uncertainty δ_xy_ that would exist for experimental data. This is a simplification, given that the dynamic localization uncertainty could very well be variable in a concrete experiment, especially given the dynamic nature of the measurements (16,26). We verified the accuracy of our analysis pipeline for analysing experimental data in terms of a well-defined set of easily interpreted physical parameters (P_hop_, D_S_, L_S_, and δ_r S_). The ranges of values for the D_S_, L_S_, and δ_r S_ parameters, explored in the simulations described below, were chosen to be comparable with the magnitudes of the equivalent parameters in the experimental datasets. The parameter P_Hop,_ on the other hand, was varied in a collection of values between 0 and 1. The trajectory length (500 localizations) and sampling rate (2kHz) of the simulated data were chosen to match those of the experimental iSCAT data. As it can be seen in Supplemental Figure S4, a very strong agreement can be reached between the ensemble average D_app_(t­_n_) curves emerging from experimental datasets and simulated trajectories, when the simulation parameters are selected accurately. Using these simulated trajectories, we were also able to explore the suitability and accuracy of our analysis pipeline on a single trajectory basis, thereby also potentially unlocking a more complete view of the heterogeneity in the observed diffusion amongst single molecules.

We then applied our quantitative ensemble average and single trajectory analysis pipelines to the simulated trajectories at analysis time ranges 0.5ms ≤ t_n_≤ T with T=5ms, 10ms, 25ms, 50ms, 75ms, and 100ms. The results of the ensemble average analysis are shown in the main text, with additional details in Supplemental Tables S6 and S7.

**Supplemental Figures and Tables referenced in the main text**

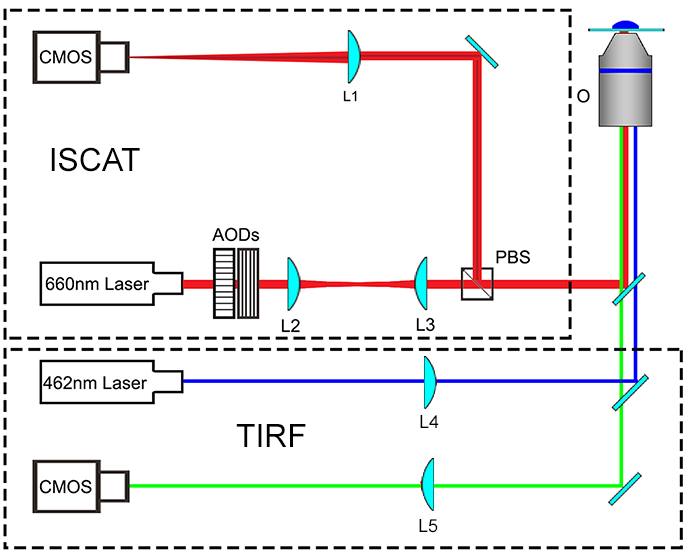

**Supplemental Figure S1 – Basic scheme of the ISCAT microscopy setup employed for the experiments.** The beam from a 660nm laser diode is scanned in the x and y directions by two Acousto-Optic deflectors (AODs, Gooch & Housego and AA Opto-Electronics). The scanning is then relayed by a telecentric system (L2, L3) to the back focal plane of the objective (O, Olympus PlanApo, 60x 1.42 NA oil immersion), mounted in inverted geometry. The component of the incident light backscattered by the sample and the reflection of the beam caused by the glass-sample interface are collected by the same objective. The polarization of the beam is adjusted so that the returning beams are then reflected by a Polarizing Beam Splitter (PBS) on the imaging camera (CMOS, Photon Focus MV-D1024-160-CL-8). The ISCAT image is formed on the camera using a 1000mm tube lens (L1, Thorlabs), so that the effective pixel size on the sensor is 31.8nm. Simultaneous fluorescence imaging was achieved by focusing the light (L4) from a 462nm laser diode in TIRF Mode with the use of a moving mirror. The fluorescence signal was detected by a second CMOS camera (FLIR Machine Vision, Grasshopper 3), after focusing with a 500mm lens (L5).

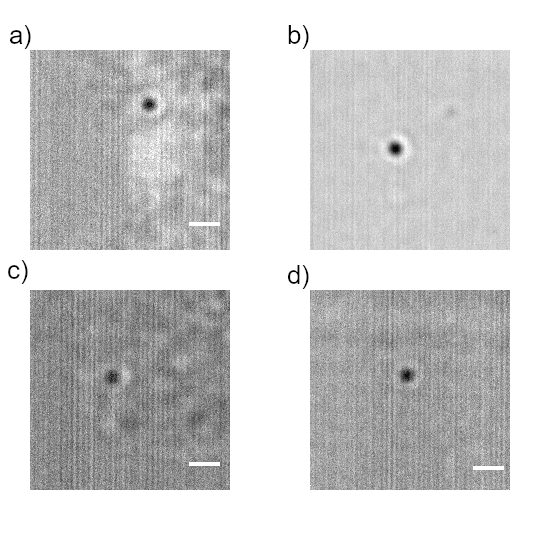

**Supplemental Figure S2 – Evaluation of scattering contrast of the gold nanoparticle tags in different environment.** a) and b): Streptavidin-coated 40nm-diameter gold nanoparticle, respectively, on a PTK2 cell surface and on immobilized on a glass substrate. c) and d): Streptavidin-coated 20nm-diameter gold nanoparticle, respectively, on a PTK2 cell surface and on immobilized on a glass substrate. On the ISCAT images detected on cell surfaces, we applied a median filter and temporal average to enhance the contrast. On the nanoparticles immobilized on glass, only a median filter was applied.

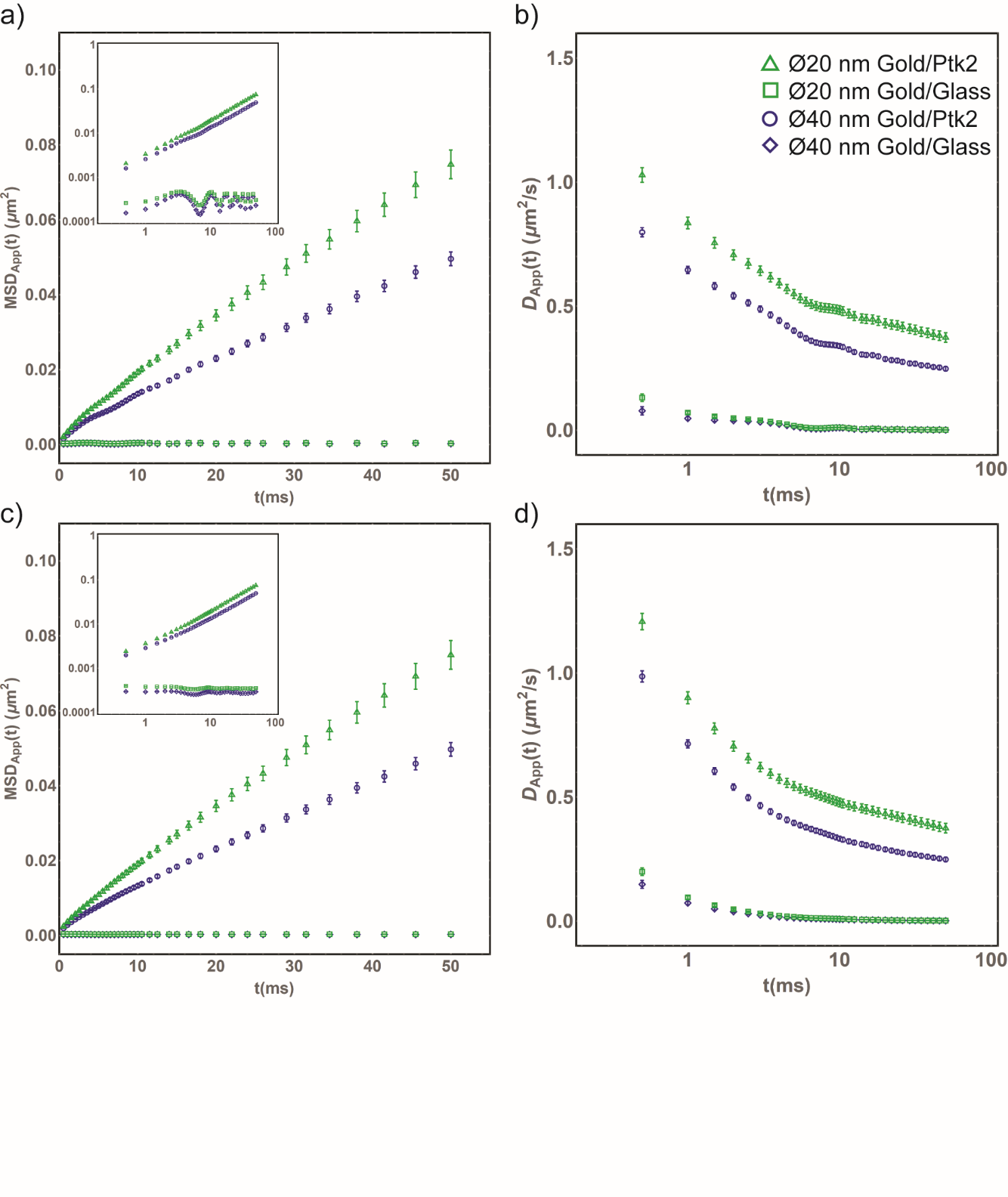

**Supplemental Figure S3** – **Comparison between vibration corrected and original Mean Squared Displacement and Apparent Diffusion Coefficient curves.** a) Ensemble average of all experimental Mean Squared Displacement curves obtained from the experimental trajectories, before vibration correction. In the insert, the same data but in a log-log representation, to further highlight the presence of environmental vibrations. b) Ensemble average of all Apparent Diffusion curves obtained from the experimental trajectories, before vibration correction. c) Same data as in a), after median filtering out in Fourier space out the frequencies in Supplemental Table S1. In the insert, the same data but in a log-log representation. d) Same data as in b), after median filtering out in Fourier space the frequencies in Supplemental Table S1

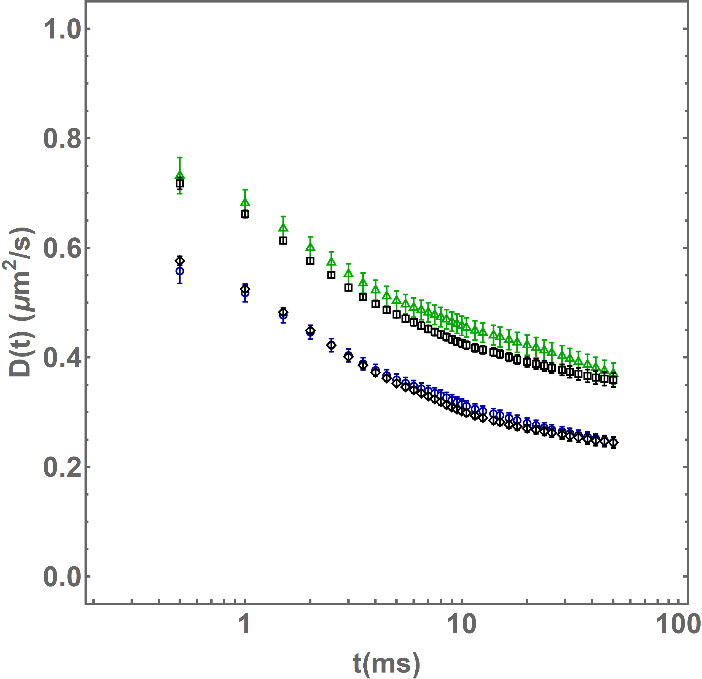

**Supplemental Figure S4 – Comparison between experimental and matching simulations.** We report the localization uncertainty-corrected, ensemble average D(t_n_) curves obtained from the datasets of experimental data (Ø20 nm gold-tagged DSPE lipids on PTK2 cell membranes in green triangles, Ø40 nm gold-tagged DSPE lipids on PTK2 cell membranes in blue squares), and the two datasets of simulated diffusion on a compartmentalized environment computed to closely match them. The parameters for the simulations were: P_Hop_=0.06, D=1.1 μm^2^/s, L=120 nm, and δ_r_=16 nm (green squares, matching the Ø20 nm gold-tagged DSPE lipids data) and P_Hop_=0.04, D=0.8 μm^2^/s, L=120 nm, and δ_r_=16 nm (black circles, matching the Ø40 nm gold-tagged DSPE lipids data)

| Sample | Frequency (±st.d.) (Hz) | Equivalent rpm (±st.d.) | RMS (±st.d.) (nm) |
| --- | --- | --- | --- |
| DSPE -PEG2000-biotin / sAv-Au (Ø20nm) on Ptk2 cells | 143±3 | 8600±200 | 7.8 ± 2.7 |
| DSPE-PEG2000-Biotin / sAv-Au (Ø40nm) on Ptk2 cells | 143±3 | 8600±200 | 7.5 ± 3.1 |
| sAv-Au (Ø20nm) immobilized on glass | 143±3 | 8600±150 | 4.2 ± 1.4 |
| sAv-Au (Ø40nm) immobilized on glass | 143±2 | 8600±130 | 4.4 ± 1.4 |

**Supplemental Table S1 – Vibration correction summary statistics.** Herein are reported the values of the frequencies, in Fourier space, that were identified to be filtered out to correct for environmental vibrations. The Equivalent RPM column is present for reference to commonly used fans for electronic equipment that might have originated the artefact.

**Supplemental Table S2 – Parameters for the model fit to the ensemble average curve of the Apparent Diffusion Coefficient data for all trajectories of Ø20nm diameter gold nanoparticles-tagged DSPE lipids on PtK2 cells.** Although all the models from Table 2 (Main text) were fit, we include here only the models for which the Relative Likelihood is larger than 0.5.

| Model | BIC | Fit Quality (p<0.05) | Relative Likelihood | D_μ_ (±S.E.)  (μm^2^/s) | D_M_ (±S.E.)  (μm^2^/s) | L (±S.E.)  (nm) | | τ_Conf_  (ms) | δ_r_ (±S.E.) (nm) | | S_Conf_ (±S.E.) |
| --- | --- | --- | --- | --- | --- | --- | --- | --- | --- | --- | --- |
| Analysis Time Range: 0.5 ≤ n δt ≤ 5 ms | | | | | | | | | | | |
| Compartmentalized Model 1 (Approximate) | -129.9 | Good | 0.72 | 0.877±0.003 | 0.423±0.001 | 99±0.4 | 5.8±0.1 | | 13.6±0.1 | 2.07±0.01 | |
| Compartmentalized Model 2 (Exact) | -130.5 | Good | 1.00 | 0.979±0.004 | 0.423±0.001 | 95±0.3 | 5.3±0.1 | | 13.4±0.1 | 2.32±0.01 | |
| Analysis Time Range: 0.5 ≤ n δt ≤ 10 ms | | | | | | | | | | | |
| Compartmentalized Model 1 (Approximate) | -197.8 | Good | 0.92 | 0.869±0.009 | 0.421±0.001 | 100±0.8 | 5.9±0.1 | | 13.6±0.1 | 2.06±0.02 | |
| Compartmentalized Model 2 (Exact) | -198.0 | Good | 1.00 | 0.969±0.011 | 0.421±0.001 | 96±0.7 | 5.5±0.1 | | 13.5±0.1 | 2.30±0.03 | |
| Analysis Time Range: 0.5 ≤ n δt ≤ 25 ms | | | | | | | | | | | |
| Compartmentalized Model 1 (Approximate) | -241.3 | Good | 0.88 | 0.82±0.02 | 0.408±0.02 | 110±2 | 7.0±0.2 | | 14.1±0.2 | 2.00±0.04 | |
| Compartmentalized Model 2 (Exact) | -241.6 | Good | 1.00 | 0.91±0.02 | 0.408±0.02 | 100 ±2 | 6.5±0.2 | | 13.9±0.2 | 2.23±0.05 | |
| Analysis Time Range: 0.5 ≤ n δt ≤ 50 ms | | | | | | | | | | | |
| Compartmentalized Model 1 (Approximate) | -242.0 | Good | 0.86 | 0.78±0.02 | 0.398±0.004 | 110±3.0 | 8.1±0.4 | | 14.4±0.2 | 1.96±0.05 | |
| Compartmentalized Model 2 (Exact) | -242.3 | Good | 1.00 | 0.87±0.02 | 0.398±0.004 | 110 ±3.0 | 7.5±0.4 | | 14.2±0.2 | 2.18±0.06 | |
| Analysis Time Range: 0.5 ≤ n δt ≤ 75 ms | | | | | | | | | | | |
| Compartmentalized Model 1 (Approximate) | -235.9 | Good | 0.84 | 0.77±0.02 | 0.393±0.005 | 120±4 | 8.8±0.6 | | 14.5±0.2 | 1.95±0.06 | |
| Compartmentalized Model 2 (Exact) | -236.3 | Good | 1.00 | 0.85±0.02 | 0.393±0.005 | 110±4 | 8.1±0.5 | | 14.4±0.2 | 2.16±0.07 | |
| Analysis Time Range: 0.5 ≤ n δt ≤ 100 ms | | | | | | | | | | | |
| Compartmentalized Model 1 (Approximate) | -237.9 | Good | 0.82 | 0.759±0.020 | 0.390±0.005 | 120±3.9 | 9.2±0.6 | | 14.6±0.2 | 1.94±0.06 | |
| Compartmentalized Model 2 (Exact) | -238.3 | Good | 1.00 | 0.84±0.02 | 0.390±0.005 | 120±3.8 | 8.5±0.6 | | 14.4±0.2 | 2.15±0.07 | |

**Supplemental Table S3. Parameters for the model fit to the ensemble average curve of the Apparent Diffusion Coefficient data for all trajectories of Ø40nm diameter gold nanoparticles-tagged DSPE lipids on PtK2 cells.** Although all the models from Table 2 (Main text) were fit, we include here only the models for which the Relative Likelihood is larger than 0.5.

| Model | BIC | Fit Quality (p<0.05) | Relative Likelihood | D_μ_ (±S.E.)  (μm^2^/s) | D_M_ (±S.E.)  (μm^2^/s) | | L (±S.E.)  (nm) | | τ_Conf_  (ms) | | δ_r_ (±S.E.) (nm) | | S_Conf_ |
| --- | --- | --- | --- | --- | --- | --- | --- | --- | --- | --- | --- | --- | --- |
| Analysis Time Range: 0.5 ≤ n δt ≤ 5 ms | | | | | | | | | | | | | |
| Compartmentalized Model 1 (Approximate) | -120.5 | Good | 0.78 | 0.695±0.005 | 0.286±0.001 | 89±0.5 | | 6.9±0.1 | | 12.7±0.0 | | 2.43±0.02 | |
| Compartmentalized Model 2 (Exact) | -121.0 | Good | 1.00 | 0.787±0.006 | 0.286±0.001 | 86±0.5 | | 6.5±0.1 | | 12.6±0.0 | | 2.75±0.02 | |
| Analysis Time Range: 0.5 ≤ n δt ≤ 10 ms | | | | | | | | | | | | | |
| Compartmentalized Model 1 (Approximate) | -189.3 | Good | 0.89 | 0.675±0.010 | 0.281±0.002 | 91±0.9 | | 7.4±0.2 | | 12.9±0.1 | | 2.41±0.04 | |
| Compartmentalized Model 2 (Exact) | -189.5 | Good | 1.00 | 0.764±0.012 | 0.281±0.002 | 88±0.9 | | 7.0±0.2 | | 12.8±0.1 | | 2.72±0.05 | |
| Analysis Time Range: 0.5 ≤ n δt ≤ 25 ms | | | | | | | | | | | | | |
| Compartmentalized Model 1 (Approximate) | -239.6 | Good | 0.82 | 0.621±0.014 | 0.265±0.003 | 100±1.9 | | 9.4±0.5 | | 13.4±0.2 | | 2.35±0.06 | |
| Compartmentalized Model 2 (Exact) | -240.0 | Good | 1.00 | 0.700±0.017 | 0.265±0.003 | 97±1.8 | | 8.8±0.4 | | 13.2±0.2 | | 2.65±0.07 | |
| Analysis Time Range: 0.5 ≤ n δt ≤ 50 ms | | | | | | | | | | | | | |
| Compartmentalized Model 1 (Approximate) | -258.3 | Good | 0.76 | 0.597±0.014 | 0.257±0.003 | 110±2.4 | | 11±0.6 | | 13.7±0.2 | | 2.32±0.06 | |
| Compartmentalized Model 2 (Exact) | -258.8 | Good | 1.00 | 0.671±0.017 | 0.256±0.003 | 100 ±2.3 | | 10±0.6 | | 13.5±0.2 | | 2.62±0.07 | |
| Analysis Time Range: 0.5 ≤ n δt ≤ 75 ms | | | | | | | | | | | | | |
| Compartmentalized Model 1 (Approximate) | -258.2 | Good | 0.73 | 0.586±0.014 | 0.253±0.004 | 110±2.7 | | 12±0.8 | | 13.8±0.2 | | 2.32±0.07 | |
| Compartmentalized Model 2 (Exact) | -258.8 | Good | 1.00 | 0.658±0.017 | 0.252±0.004 | 100±2.6 | | 11±0.7 | | 13.6±0.2 | | 2.61±0.08 | |
| Analysis Time Range: 0.5 ≤ n δt ≤ 100 ms | | | | | | | | | | | | | |
| Compartmentalized Model 1 (Approximate) | -263.6 | Good | 0.71 | 0.581±0.014 | 0.251±0.004 | 110±2.8 | | 12±0.8 | | 13.8±0.2 | | 2.32±0.07 | |
| Compartmentalized Model 2 (Exact) | -264.3 | Good | 1.00 | 0.652±0.017 | 0.250±0.004 | 110±2.7 | | 11±0.8 | | 13.6±0.2 | | 2.61±0.08 | |

**Supplemental Table S4. Parameters for the model fit to the ensemble average curve of the Apparent Diffusion Coefficient data for all trajectories of Ø20nm diameter gold nanoparticles immobilized on glass substrate.** Although all the models from Table 2 (Main text) were fit, we include here only the models for which the Relative Likelihood is larger than 0.5.

| Model | BIC | Relative Likelihood  All Models | Fit Quality (p<0.05) | D_μ_ (±S.E.)  (μm^2^/s) | D_M_ (±S.E.)  (μm^2^/s) | | L (±S.E.)  (nm) | | τ_Conf_ (±S.E.)  (ms) | | δ_r_ (±S.E.) (nm) | | S_Conf_ (±S.E.) |
| --- | --- | --- | --- | --- | --- | --- | --- | --- | --- | --- | --- | --- | --- |
| Analysis Time Range: 0.5 ≤ n δt ≤ 5 ms | | | | | | | | | | | | | |
| Localization Uncertainty | -88.8 | 6.7x10^-4^ | Good |  |  |  | |  | | 9.3±0.1 | |  | |
| Analysis Time Range: 0.5 ≤ n δt ≤ 10 ms | | | | | | | | | | | | | |
| Localization Uncertainty | -194.4 | 3.1x10^-6^ | Good |  |  |  | |  | | 9.3±0.1 | |  | |
| Analysis Time Range: 0.5 ≤ n δt ≤ 25 ms | | | | | | | | | | | | | |
| Localization Uncertainty | -328.8 | 1.3x10^-8^ | Good |  |  |  | |  | | 9.3±0.1 | |  | |
| Analysis Time Range: 0.5 ≤ n δt ≤ 50 ms | | | | | | | | | | | | | |
| Localization Uncertainty | -398.1 | 6.6x10^-10^ | Good |  |  |  | |  | | 9.3±0.1 | |  | |
| Analysis Time Range: 0.5 ≤ n δt ≤ 75 ms | | | | | | | | | | | | | |
| Localization Uncertainty | -421.4 | 3.3x10^-10^ | Good |  |  |  | |  | | 9.3±0.1 | |  | |
| Analysis Time Range: 0.5 ≤ n δt ≤ 100 ms | | | | | | | | | | | | | |
| Localization Uncertainty | -445.0 | 9.2x10^-10^ | Good |  |  |  | |  | | 9.3±0.1 | |  | |

**Supplemental Table S5. Parameters for the model fit to the ensemble average curve of the Apparent Diffusion Coefficient data for all trajectories of Ø40nm diameter gold nanoparticles immobilized on glass substrate.** Although all the models from Table 2 (Main text) were fit, we include here only the models for which the Relative Likelihood is larger than 0.5.

| Model | BIC | Relative Likelihood  All Models | Fit Quality (p<0.05) | D_μ_ (±S.E.)  (μm^2^/s) | D_M_ (±S.E.)  (μm^2^/s) | | L (±S.E.)  (nm) | | τ_Conf_ (±S.E.)  (ms) | | δ_r_ (±S.E.) (nm) | | S_Conf_ (±S.E.) |
| --- | --- | --- | --- | --- | --- | --- | --- | --- | --- | --- | --- | --- | --- |
| Analysis Time Range: 0.5 ≤ n δt ≤ 5 ms | | | | | | | | | | | | | |
| Localization Uncertainty | -85.8 | 5.8x10^-10^ | Good |  |  |  | |  | | 8.1±0.1 | |  | |
| Free | -89.4 | 3.6x10^-9^ | Good |  | 0.003±0.001 |  | |  | | 7.9±0.1 | |  | |
| Analysis Time Range: 0.5 ≤ n δt ≤ 10 ms | | | | | | | | | | | | | |
| Localization Uncertainty | -187.5 | 1.3x10^-6^ | Good |  |  |  | |  | | 8.1±0.1 | |  | |
| Free | -190.1 | 4.8x10^-6^ | Good |  | 0.001±0.001 |  | |  | | 8.0±0.1 | |  | |
| Analysis Time Range: 0.5 ≤ n δt ≤ 25 ms | | | | | | | | | | | | | |
| Localization Uncertainty | -316.8 | 7.3x10^-11^ | Good |  |  |  | |  | | 8.1±0.1 | |  | |
| Free | -320.9 | 5.6x10^-10^ | Good |  | 0.001±0.001 |  | |  | | 8.0±0.1 | |  | |
| Analysis Time Range: 0.5 ≤ n δt ≤ 50 ms | | | | | | | | | | | | | |
| Localization Uncertainty | -383.7 | 5.7x10^-13^ | Good |  |  |  | |  | | 8.1±0.1 | |  | |
| Free | -387.2 | 3.3x10^-12^ | Good |  | 0.001±0.0005 |  | |  | | 8.0±0.1 | |  | |
| Analysis Time Range: 0.5 ≤ n δt ≤ 75 ms | | | | | | | | | | | | | |
| Localization Uncertainty | -406.5 | 1.2x10^-13^ | Good |  |  |  | |  | | 8.1±0.1 | |  | |
| Free | -409.3 | 5.0x10^-13^ | Good |  | 0.001±0.0005 |  | |  | | 8.0±0.1 | |  | |
| Analysis Time Range: 0.5 ≤ n δt ≤ 100 ms | | | | | | | | | | | | | |
| Localization Uncertainty | -428.9 | 2.2x10^-14^ | Good |  |  |  | |  | | 8.1±0.1 | |  | |
| Free | -431.8 | 9.4x10^-14^ | Good |  | 0.001±0.0005 |  | |  | | 8.0±0.1 | |  | |

**Supplemental Table S6. Model Fit parameters for simulated trajectories matching the Ø20nm gold tagged DSPE lipids diffusing on PTK2 cell membranes.** A set of 100 trajectories, 500 localizations long at 2kHz sampling rates were simulated, and their D_app_(t_n_) curves averaged together. The resulting ensemble average curves were Simulation parameters are: P_hop_ = 0.06, D = 1.0 μm^2^/s, L = 120nm, δ_r_ = 8nm

| Model | BIC | Fit Quality (p<0.05) | Relative Likelihood | D_μ_ (±S.E.)  (μm^2^/s) | D_M_ (±S.E.)  (μm^2^/s) | L (±S.E.)  (nm) | τ_Conf_  (ms) | δ_r_ (±S.E.) (nm) | S_Conf_ (±S.E.) |
| --- | --- | --- | --- | --- | --- | --- | --- | --- | --- |
| Analysis Time Range: 0.5 ≤ n δt ≤ 5 ms | | | | | | | | | |
| Compartmentalized Model 1 (Approximate) | -109.71 | Good | 0.987 | 0.865 ± 0.01 | 0.397 ± 0.002 | 97.4 ± 0.9 | 6.0 ± 0.2 | 13.1 ± 0.1 | 2.18 ± 0.03 |
| Compartmentalized Model 2 (Exact) | -109.74 | Good | 1 | 0.974 ± 0.01 | 0.397 ± 0.002 | 93.8 ± 0.9 | 5.5 ± 0.1 | 12.9 ± 0.1 | 2.45± 0.03 |
| Analysis Time Range: 0.5 ≤ n δt ≤ 10 ms | | | | | | | | | |
| Compartmentalized Model 1 (Approximate) | -179.398 | Good | 0.964 | 0.825 ± 0.01 | 0.386 ± 0.002 | 102 ± 1 | 6.8 ± 0.2 | 13.5 ± 0.1 | 2.14 ± 0.04 |
| Compartmentalized Model 2 (Exact) | -179.471 | Good | 1 | 0.927 ± 0.02 | 0.386 ± 0.002 | 98 ± 1 | 6.3 ± 0.2 | 13.3 ± 0.1 | 2.4 ± 0.04 |
| Analysis Time Range: 0.5 ≤ n δt ≤ 25 ms | | | | | | | | | |
| Compartmentalized Model 1 (Approximate) | -249.565 | Good | 0.916 | 0.787 ± 0.01 | 0.375 ± 0.002 | 107 ± 1 | 7.7 ± 0.2 | 13.9 ± 0.2 | 2.10 ± 0.04 |
| Compartmentalized Model 2 (Exact) | -249.74 | Good | 1 | 0.882 ± 0.02 | 0.375 ± 0.002 | 104 ± 1 | 7.2 ± 0.1 | 13.7 ± 0.2 | 2.35 ± 0.05 |
| Analysis Time Range: 0.5 ≤ n δt ≤ 50 ms | | | | | | | | | |
| Compartmentalized Model 1 (Approximate) | -271.32 | Good | 0.895 | 0.77 ± 0.01 | 0.371 ± 0.003 | 110 ± 2 | 8.2 ± 0.3 | 14.1 ± 0.2 | 2.08 ± 0.04 |
| Compartmentalized Model 2 (Exact) | -271.54 | Good | 1 | 0.86 ± 0.02 | 0.371 ± 0.003 | 106 ± 2 | 7.6 ± 0.3 | 13.8 ± 0.2 | 2.33 ± 0.05 |
| Analysis Time Range: 0.5 ≤ n δt ≤ 75 ms | | | | | | | | | |
| Compartmentalized Model 1 (Approximate) | -273.365 | Good | 0.889 | 0.77 ± 0.01 | 0.369 ± 0.003 | 111 ± 2 | 8.4 ± 0.3 | 14.1 ± 0.2 | 2.08 ± 0.04 |
| Compartmentalized Model 2 (Exact) | -273.602 | Good | 1 | 0.86 ± 0.02 | 0.369 ± 0.003 | 107 ± 2 | 7.7 ± 0.3 | 13.9 ± 0.2 | 2.32 ± 0.05 |
| Analysis Time Range: 0.5 ≤ n δt ≤ 100 ms | | | | | | | | | |
| Compartmentalized Model 1 (Approximate) | -280.945 | Good | 0.883 | 0.77 ± 0.01 | 0.369 ± 0.003 | 111 ± 2 | 8.4 ± 0.3 | 14.2 ± 0.2 | 2.08 ± 0.04 |
| Compartmentalized Model 2 (Exact) | -281.192 | Good | 1 | 0.86 ± 0.02 | 0.369 ± 0.003 | 107 ± 2 | 7.7 ± 0.3 | 13.9 ± 0.2 | 2.32 ± 0.05 |

**Supplemental Table S7. Ensemble Average Analysis Results for simulated trajectories (Parameters: P_hop_ = 0.04, D = 0.8 μm^2^/s, L = 120nm, δ_r_ = 8nm**)

| Model | BIC | Fit Quality (p<0.05) | Relative Likelihood | D_μ_ (±S.E.)  (μm^2^/s) | D_M_ (±S.E.)  (μm^2^/s) | L (±S.E.)  (nm) | τ_Conf_  (ms) | δ_r_ (±S.E.) (nm) | S_Conf_ (±S.E.) |
| --- | --- | --- | --- | --- | --- | --- | --- | --- | --- |
| Analysis Time Range: 0.5 ≤ n δt ≤ 5 ms | | | | | | | | | |
| Compartmentalized Model 1 (Approximate) | -116.202 | Good | 1 | 0.679 ± 0.006 | 0.267 ± 0.002 | 93.4 ± 0.8 | 8.16 ± 0.3 | 12.5 ± 0.07 | 2.54 ± 0.03 |
| Compartmentalized Model 2 (Exact) | -116.043 | Good | 0.923 | 0.774 ± 0.007 | 0.267 ± 0.002 | 90.6 ± 0.7 | 7.7 ± 0.2 | 12.2 ± 0.07 | 2.9 ± 0.03 |
| Analysis Time Range: 0.5 ≤ n δt ≤ 10 ms | | | | | | | | | |
| Compartmentalized Model 1 (Approximate) | -183.98 | Good | 0.97 | 0.66 ± 0.01 | 0.261 ± 0.002 | 96 ± 1 | 8.9 ± 0.3 | 12.7 ± 0.1 | 2.53 ± 0.04 |
| Compartmentalized Model 2 (Exact) | -184.07 | Good | 1 | 0.75 ± 0.01 | 0.261 ± 0.002 | 93 ± 1 | 8.3 ± 0.3 | 12.4 ± 0.1 | 2.9 ± 0.05 |
| Analysis Time Range: 0.5 ≤ n δt ≤ 25 ms | | | | | | | | | |
| Compartmentalized Model 1 (Approximate) | -261.46 | Good | 0.88 | 0.631 ± 0.01 | 0.251 ± 0.002 | 101 ± 1 | 10.1 ± 0.4 | 13.0 ± 0.1 | 2.51 ± 0.04 |
| Compartmentalized Model 2 (Exact) | -261.7 | Good | 1 | 0.72 ± 0.01 | 0.251 ± 0.002 | 98 ± 1 | 9.5 ± 0.3 | 12.8 ± 0.1 | 2.86 ± 0.05 |
| Analysis Time Range: 0.5 ≤ n δt ≤ 50 ms | | | | | | | | | |
| Compartmentalized Model 1 (Approximate) | -293.17 | Good | 0.85 | 0.62 ± 0.01 | 0.248 ± 0.002 | 102 ± 1 | 10.5 ± 0.4 | 13.1 ± 0.1 | 2.51 ± 0.04 |
| Compartmentalized Model 2 (Exact) | -293.5 | Good | 1 | 0.71 ± 0.01 | 0.248 ± 0.002 | 99 ± 1 | 9.9 ± 0.4 | 12.9 ± 0.1 | 2.85 ± 0.05 |
| Analysis Time Range: 0.5 ≤ n δt ≤ 75 ms | | | | | | | | | |
| Compartmentalized Model 1 (Approximate) | -295 | Good | 0.85 | 0.62 ± 0.01 | 0.247 ± 0.002 | 103 ± 1 | 10.7 ± 0.4 | 13.1 ± 0.1 | 2.51 ± 0.05 |
| Compartmentalized Model 2 (Exact) | -295.3 | Good | 1 | 0.70 ± 0.01 | 0.247 ± 0.002 | 100 ± 1 | 10.1 ± 0.4 | 12.9 ± 0.1 | 2.85 ± 0.06 |
| Analysis Time Range: 0.5 ≤ n δt ≤ 100 ms | | | | | | | | | |
| Compartmentalized Model 1 (Approximate) | -299.3 | Good | 0.845 | 0.62 ± 0.01 | 0.246 ± 0.002 | 103 ± 1 | 10.9 ± 0.5 | 13.2 ± 0.1 | 2.5 ± 0.05 |
| Compartmentalized Model 2 (Exact) | -299.63 | Good | 1 | 0.70 ± 0.01 | 0.246 ± 0.002 | 100 ± 1 | 10.2 ± 0.4 | 12.9 ± 0.2 | 2.85 ± 0.06 |
